# Side-necked turtle genomes reveal chromosomal dynamics, skeletal innovation and cancer resistance

**DOI:** 10.64898/2026.03.05.709825

**Authors:** Leon Hilgers, Michail Rovatsos, Dimitrios-Georgios Kontopoulos, Thomas Brown, Thomas Hickler, Brian Huntley, Martin Pippel, Chetan Munegowda, Tobias Müller, Alexis-Walid Ahmed, Annemarie Laas, Peter Praschag, Joana Damas, Sylke Winkler, Harris Lewin, Eugene Myers, Uwe Fritz, Michael Hiller

## Abstract

Turtles exhibit a highly derived body plan, exceptional longevity, cancer resistance, and striking diversity in karyotypes and sex determination systems. However, the genomic basis of these innovations remains unresolved, largely because reference genomes were lacking for one of two extant turtle clades, the side-necked turtles (Pleurodira). As part of the Vertebrate Genomes Project, we generated seven reference-quality Pleurodira genomes and reconstructed the most comprehensive genome-wide turtle phylogeny. Combining demographic inference with historical climate and biome reconstructions indicates that ancient climate fluctuations shaped long-term population dynamics, while recent declines mainly reflect population structure. Ancestral genome reconstructions reveal that rare bursts of chromosome fusions and fissions, likely facilitated by repetitive elements, drove turtle karyotype diversity. By identifying sex chromosomes and tracing their evolutionary history, we resolve a long-standing debate and demonstrate a single origin of genetic sex determination in Chelidae on a microchromosome over 80 million years ago. Contrary to previous hypotheses, we find no evidence of coevolution between genetic sex determination and chromosome number in turtles. Comparative genomic analyses further identify gene losses and signatures of adaptive evolution associated with key turtle traits. Gene losses causing disproportionate dwarfism phenotypes may have contributed to skeletal adaptations underlying the compact turtle body plan. In addition, gene losses and adaptive changes in stress response and tumor suppressor pathways likely enhance oxidative stress tolerance and cancer resistance. Together, these findings illuminate turtle genome evolution, revealing chromosomal dynamics, sex chromosome evolution, molecular insights into skeletal innovation and cancer resistance, and implicate gene losses as a recurrent contributor to evolutionary novelty.

## Introduction

Understanding the drivers of genome evolution and illuminating how molecular change gives rise to adaptation and evolutionary innovation are central goals of evolutionary biology. Here, turtles offer a unique opportunity: referred to as the ultimate hopeful monsters due to their radically different body plan ^1,2^, turtles originated ∼250 million years ago when climate changes caused mass extinctions, the most extreme of which (Permian-Triassic mass extinction) pushed >85% of all species into extinction ^3–5^. During this time, stem-turtles, i.e., the extinct ancestors of modern turtles, reconfigured the tetrapod skeleton, which gave rise to the unique turtle body plan including one of the most iconic evolutionary novelties: the turtle shell ^6,7^. While turtles were much more species rich in the past ^8^, extant turtles comprise 366 species in two clades, the side-necked turtles (Pleurodira, 95 species) and the hidden necked turtles (Cryptodira, 271 species)^9,10^.

Turtles also exhibit physiological adaptations including remarkable longevity with negligible ageing, natural cancer resistance and pronounced hypoxia tolerance ^11–14^. Even the smallest turtle species surpass 20 years in lifespan, while some giants, such as the Galápagos tortoise and Aldabra tortoise, live for more than 150 years ^13^. Analyses of giant tortoise genomes found signatures of positive selection and duplication of genes involved in genome maintenance, tumor suppression and ageing ^15^. However, not only giant tortoises, but turtles in general exhibit remarkably low cancer prevalence (∼ 0% to 0.78%) compared to other vertebrates including birds (1% in doves and pigeons to 17% in landfowl) and mammals (3% in bats to 17% in carnivores) ^16^. This cancer resistance of turtles may be linked to their exceptional oxidative stress tolerance, which requires adaptations to avoid or effectively repair DNA damage ^17^. The earliest stem-turtles were likely terrestrial with a fossorial lifestyle ^18^, but ancestors of all living turtles secondarily adapted to an aquatic life ^19,20^. Thus, already early in their evolutionary history turtles adapted to hypoxic conditions and the increased oxidative stress associated with hypoxia and reoxygenation ^21^. In line with this idea, previous work indicates that turtles are resilient to oxidative stress ^11,17,22–24^ and extant turtle species are among the most hypoxia-tolerant vertebrates on the planet ^21^. However, while a few studies have provided initial insights into longevity, cancer resistance and hypoxia tolerance in turtles ^11,15,22,25^, the genomic basis of these traits remains largely unknown.

In addition to morphological and physiological adaptations, turtles exhibit a fascinating karyotype diversity ranging from n=13 to n=34 ^26^. Chromosome-level genome assemblies make it possible to reveal chromosome fission and fusion events and thus reconstruct the processes underlying karyotype diversity. Understanding the mechanisms and drivers of chromosome evolution is important because they change evolutionary trajectories by altering recombination landscapes and gene regulation, and by creating reproductive barriers that promote speciation ^27,28^.

Turtles also vary in their sex determination. The majority of extant species have temperature-dependent sex determination (TSD) ^29^, which is considered their ancestral sex determination system ^26,30,31^. However, at least five lineages of turtles independently evolved genotypic sex determination (GSD), either under male (XX/XY) or female (ZW) heterogamety (reviewed in ^26,30^). Interestingly, researchers discovered both micro- and macro-sex chromosomes within a single clade of side-necked turtles, the Chelidae ^32–34^, sparking a debate about how these sex chromosomes evolved and whether genetic sex determination originated once, or twice independently in this clade ^33–35^.

Today, the majority of turtle species are classified under a threatened category according to the IUCN Red list (https://www.iucnredlist.org/), which makes turtles one of the most endangered major vertebrate clades ^9,36,37^. Global turtle population collapses already impair ecological roles that turtles have played for more than 200 million years across a wide range of ecosystems from deserts to the open ocean ^36^. Key drivers of turtle declines include habitat loss, overexploitation and climate change ^9,37^. Here, climate change may play a particularly devastating role for turtles that rely on temperature-dependent sex determination, because incubation temperatures are expected to change. In some cases, this may already have created maladaptive sex ratio biases ^37–40^. Thus, it is important to understand how climate- and ecosystem change will affect turtle populations and whether the impact of climate change differs between species with genetic- and temperature dependent sex determination.

Reference-quality turtle genomes can shed light on the genomic underpinnings of turtle traits such as cancer resistance and longevity, illuminate the evolution of chromosome diversity and genetic sex determination, uncover historical population dynamics, and provide important resources to improve turtle conservation. However, until recently, reference genomes were only available for one of two living turtle suborders, the hidden-neck turtles (Crypodira), and entirely lacking for the side-necked turtle (Pleurodira) suborder, which limited comparative analyses.

Here, as part of the Vertebrate Genomes Project (VGP) effort to generate reference genomes for all vertebrate species, we generated seven reference-quality side-necked turtle genomes. Using our newly generated resources, we infer population histories to explore the effects of historical climate- and biome change on turtle populations. We further identify sex chromosomes, illuminate their evolution in Chelidae and reconstruct chromosome evolution across the turtles to explore drivers of chromosome evolution.

Finally, we link gene losses and signatures of positive selection in stem-turtles to turtle-specific novelties including their unique body plan and their remarkable cancer resistance.

## Results

### Seven new reference-quality side necked turtle genomes

To fill the gap of reference genomes for side-necked turtles (Pleurodira), we provide seven new chromosome-level genome assemblies. We cover all three major clades of side-necked turtles by sequencing three species of the Chelidae (*Elseya novaeguineae, Chelodina parkeri*, and *Hydromedusa tectifera*), three species of Podocnemididae (*Peltocephalus dumerilianus, Erymnochelys madagascariensis,* and *Podocnemis unifilis*) and one species of Pelomedusidae (*Pelomedusa somalica*). *Erymnochelys madagascariensis* further represents the tetrapod with the highest EDGE score, which indicates that this species is uniquely endangered and evolutionarily distinct: it diverged from its closest living relative ∼ 80 mya ^41,42^.

For all turtles except *Elseya novaeguineae*, we assembled genomes from PacBio continuous long reads (CLR), Bionano optical maps, 10X Genomics linked reads and HiC chromatin conformation capture data using the VGP v1.6 pipeline ^43^. For *Elseya novaeguineae*, we generated PacBio high-fidelity (HiFi) and HiC data to assemble a haplotype-resolved genome ^44^. With 97.8–99.9% of the assemblies contained in chromosome-size scaffolds and contig N50 values ranging from 42 Mb to 68 Mb, our new genomes exceed assembly quality metrics proposed by the VGP ^43^ and are as good as recently published turtle genomes ^45^ (Figure 1A). Importantly, all assemblies far exceed the contiguity of the previously available Pleurodira genome assemblies except for two very recent chromosome-level Pleurodira genomes ^46^ (Figure 1A, Table S1). We annotated coding genes with TOGA ^47^, integrating annotations generated with seven reference species. To evaluate annotation completeness, we identified complete conserved Sauropsida single copy orthologs (BUSCOs, n=7,840) ^48^. Scores ranging from 99.6% to 99.9% indicate a high completeness of our annotations and hence our new reference genomes (Figure 1B), which lay the foundation for comprehensive comparative genomics and conservation genomics in turtles.

**Figure 1:**
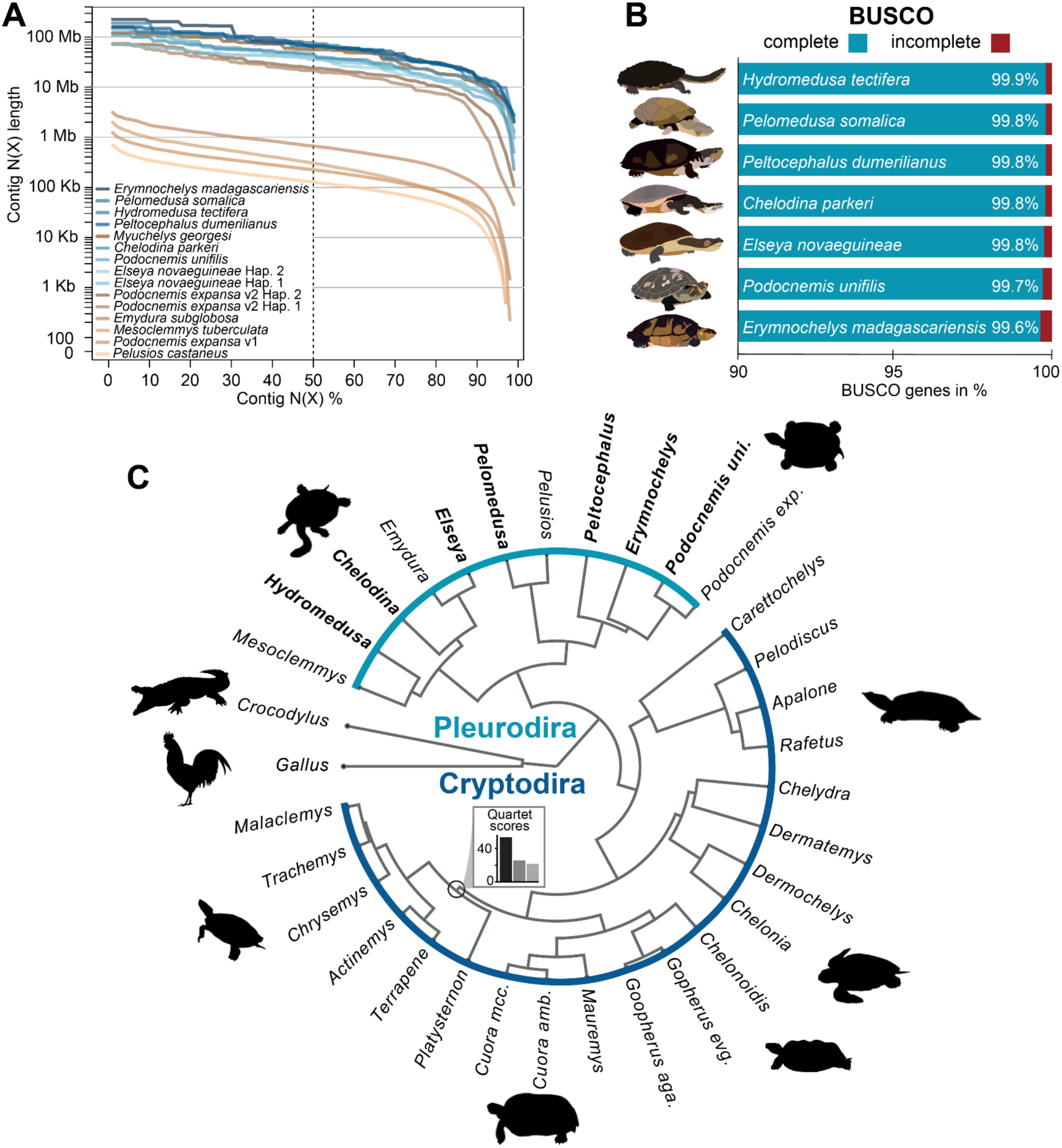
Seven new reference genomes for side-necked turtles (Pleurodira). A) Genome assembly contiguity with contig sizes on the y-axis and contig N(X) in % on the x-axis for available (brown) and our newly assembled genomes (blue). N(X) indicates that X% of the assembly consists of contigs of at least that size. B) Genome assembly completeness measured as presence and completeness of 7,840 conserved single copy orthologs (BUSCOs sauropsida odb10 database). C) Time-calibrated turtle phylogeny including our new Pleurodira genomes (bold) based on >14.000 1:1 orthologs. All nodes have posterior probabilities of 1.0 and the normalized quartet score was 0.952. The bar plot shows quartet scores for the node indicating a sister relationship of *Platysternon* and *Emydidae,* which is supported by more than twice the number of gene trees (black) compared to each of the two alternative hypotheses (grey). Chicken and crocodile were used as outgroups.

### Genome-wide phylogeny supports recent phylogenetic hypotheses

We inferred a genome-based turtle phylogeny that includes our newly sequenced species using multi-species coalescence. To this end, we generated codon-aware exon-by-exon alignments of 14,530 1:1 orthologous genes, computed gene trees, and used them to infer a species tree. This makes our dataset the largest to infer a genome-wide turtle phylogeny, increasing the number of analyzed orthologs 5.7-fold from 2,513 to 14,530 and the species number by ∼20% in comparison to the largest previously used data set^49^.

We obtained a very well-resolved phylogeny that matches previous topologies ^42,49^ with all nodes exhibiting posterior probabilities of 1.0 and a normalized quartet score of 0.952, indicating that ≥95.2% of the gene tree quartets are in agreement with the species tree (Figure 1C). With this, our phylogeny also settles a debate about the controversial sister relationship between *Platysternon* and Emydidae, as this relationship is supported by more than twice the number of gene trees in our analyses (q1 = 53%) compared to the alternative hypotheses (q2 = 25%, q3 = 22%) (Figure 1C). Similar to previous work, our data indicate very limited gene-tree species-tree discordance and provide strong evidence for a sister relationship of *Platysternon* and Emydidae ^42,49,50^.

### Geography and climate drove population sizes in the distant past

After thriving for more than 200 million years, turtles have become one of the most endangered major vertebrate clades ^9,36,37^. This recent decline of turtles raises the question of how turtles responded to global change in the past. Using our seven and four publicly-available side-necked turtle genomes, we inferred historical population size changes for 11 species from South America, Africa and Oceania (Figure 2A) with the sequentially Markovian coalescent (PSMC) method ^51^. Following our previous results, we increased the number of time windows in the recent past, which improved the resolution of PSMC and removed all artefactual peaks across our dataset ^52^. Although we observed species-specific differences in population dynamics, population sizes of all investigated species from Oceania and Africa peaked around the onset of the Mid-Pleistocene Transition (MPT) ∼1.2 – 0.8 million years ago (mya), followed by a long period of population decline (Figure 2B). This period was characterized by glacial ice sheet expansions, enhanced aridification ^53^, extreme cooling events ^54^ and a major reorganization of Asian and Australian monsoon ^53,55^, which was caused by longer and more extreme glacial-interglacial cycles (Figure 2B). In contrast to species from Oceania and Africa, the Amazonian species except for *Peltocephalus dumerilianus*, which buries itself during the dry season, exhibit increasing population sizes during and after the MPT. This finding is consistent with the MPT likely supporting the formation of larger Amazon floodplains that only retreated in the late Pleistocene ^56,57^. Our results further align with reported population declines of several freshwater Cryptodira after the MPT ^58,59^ and indicate that continent dependent differences in climatic change explain historical turtle population dynamics.

**Figure 2:**
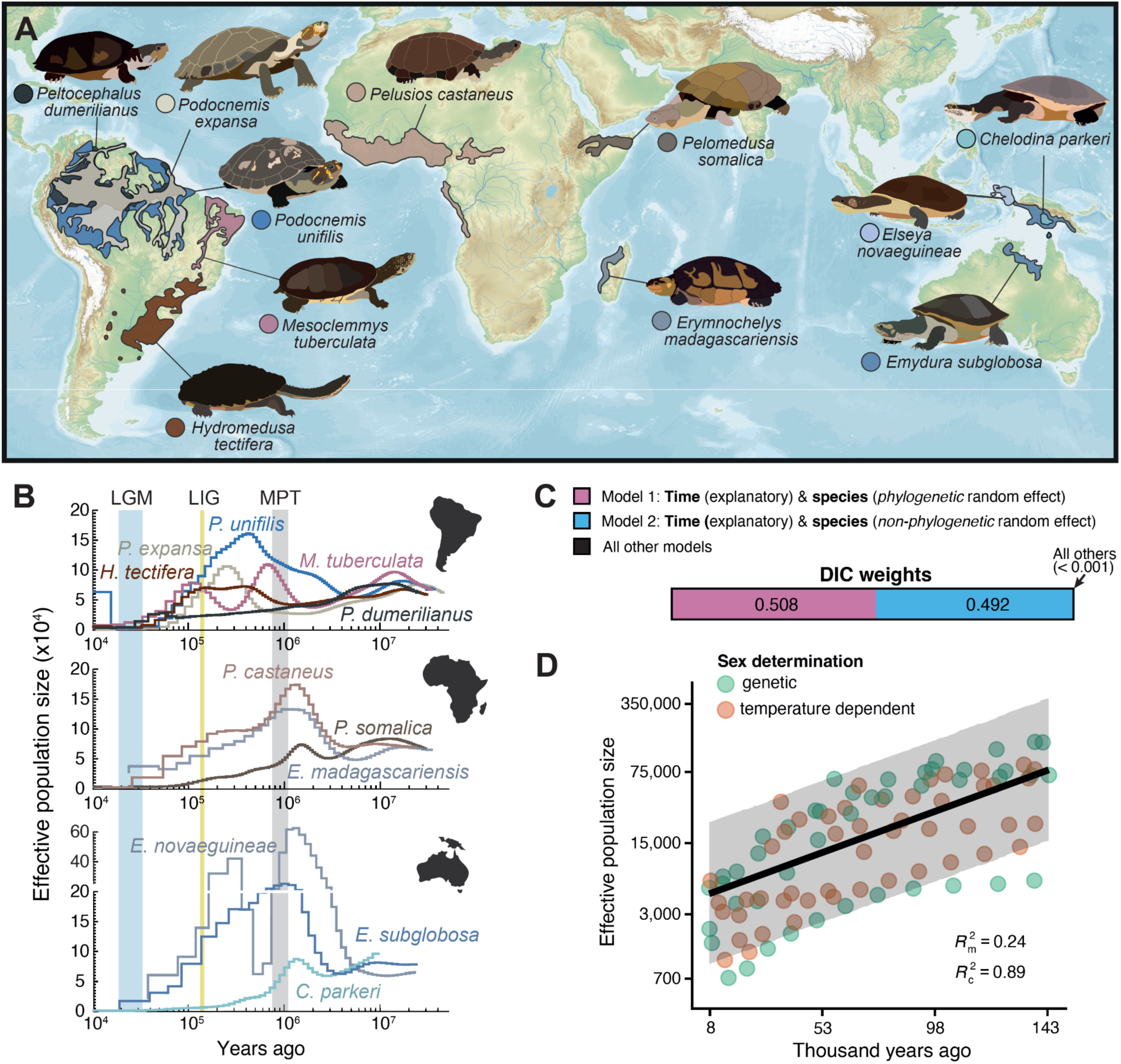
Demographic history of side necked turtles. A) Present day distribution ranges of 11 analyzed side necked turtle species. B) Population histories inferred with PSMC for species in South America (top), Africa (middle) and Oceania (bottom). Time (years ago) is shown on the logarithmic x-axis and the inferred effective population size on the y-axis. Vertical bars show three important climatic events including the last glacial maximum (LGM, blue), the last interglacial (LIG, yellow) and the Mid-Pleistocene transition (MPT, grey). C) DIC weights of 615 generalized linear mixed models to explore the effects of historical biome suitability, sex determination, mean temperature, temperature range and time, to account for declining population sizes towards the present due to population structure. D) Plot of model 1 with population size on the y-axis and time on the x-axis. In this model, time explains 24% of variance in population size (*R*^2^_m_), whereas the combination of time and a phylogenetic random effect on the intercept explains 89% of the variance (*R*^2^_c_).

### Recent population declines are likely caused by population structure

For all 11 investigated species, PSMC indicated dramatic population declines in more recent times since the last interglacial from ∼130,000–115,000 years ago (Figure 2B). To explore the cause of inferred turtle population collapses since the last interglacial, we fitted a series of generalized linear mixed models (GLMM). To this end, we collected historical climate data from the last 140,000 years ^60,61^ and scored historical biome suitability based on historical biome reconstructions ^62^ and biome compositions in present-day distribution ranges ^63^. Additionally, we included the sex determination mode in some of the tested models to explore whether the effect of past climate change differed between species with genetic and temperature-dependent sex determination. Finally, since population structure can cause population declines in recent time periods, even if population sizes remain constant ^64^, we also included time as a potential explanatory variable. Strikingly, all top-ranked models included time as the only explanatory variable. This means that after accounting for a gradual decline in population size with time, which could be caused solely by existing population structure, we do not detect a significant effect of climate, biome, or sex determination on population size.

### Identification of micro- and macro sex chromosomes in Chelidae

Chelidae evolved genetic sex determination with male heterogamety (XX/XY) from an ancestral temperature-dependent sex determination ^26,65^. However, the size of the sex chromosomes differs, with cytogenetic analyses revealing heteromorphic macro-sex chromosomes in a clade comprising *Emydura* and *Elseya* ^33,34^, but homomorphic micro-sex chromosomes in *Chelodina* ^32,34^. This raises the question whether GSD evolved multiple times independently in Chelidae. Additionally, a third clade that is sister to all other Chelidae and includes several genera such as *Phrynops*, *Mesoclemmys* and *Hydromedusa*, has a so far unknown GSD system ^34,66^.

To unravel the identity and evolution of sex chromosomes in Chelidae, we first focused on *Elseya novaeguineae*, where leveraging its haplotype-resolved assembly, we integrated results from three orthogonal analyses. First, we re-sequenced a male and a female and used Findzx ^67^ to identify sex chromosomes by localized patterns of increased heterozygosity in males (Figure 3). This analysis is based on the principle that in an XX/XY system, a non-recombining sex-linked region diverges between the X and the Y, causing a localized increase in heterozygosity in males (XY) compared to females (XX). Our data revealed that the most pronounced outlier region with increased heterozygosity in males compared to females is situated at the end of the fourth largest scaffold (Figure 4B). This result is in line with previous cytogenetic analyses showing that the chromosomal pair 4 of *Elseya* corresponds to the sex chromosomes and the Y chromosome is visible under microscope with cytogenetic stains ^34^. Microdissecting the Y chromosome of *Elseya novaeguineae* followed by sequencing and mapping the illumina reads provided further support that chromosome 4 is the sex chromosome (Figure S1).

**Figure 3:**
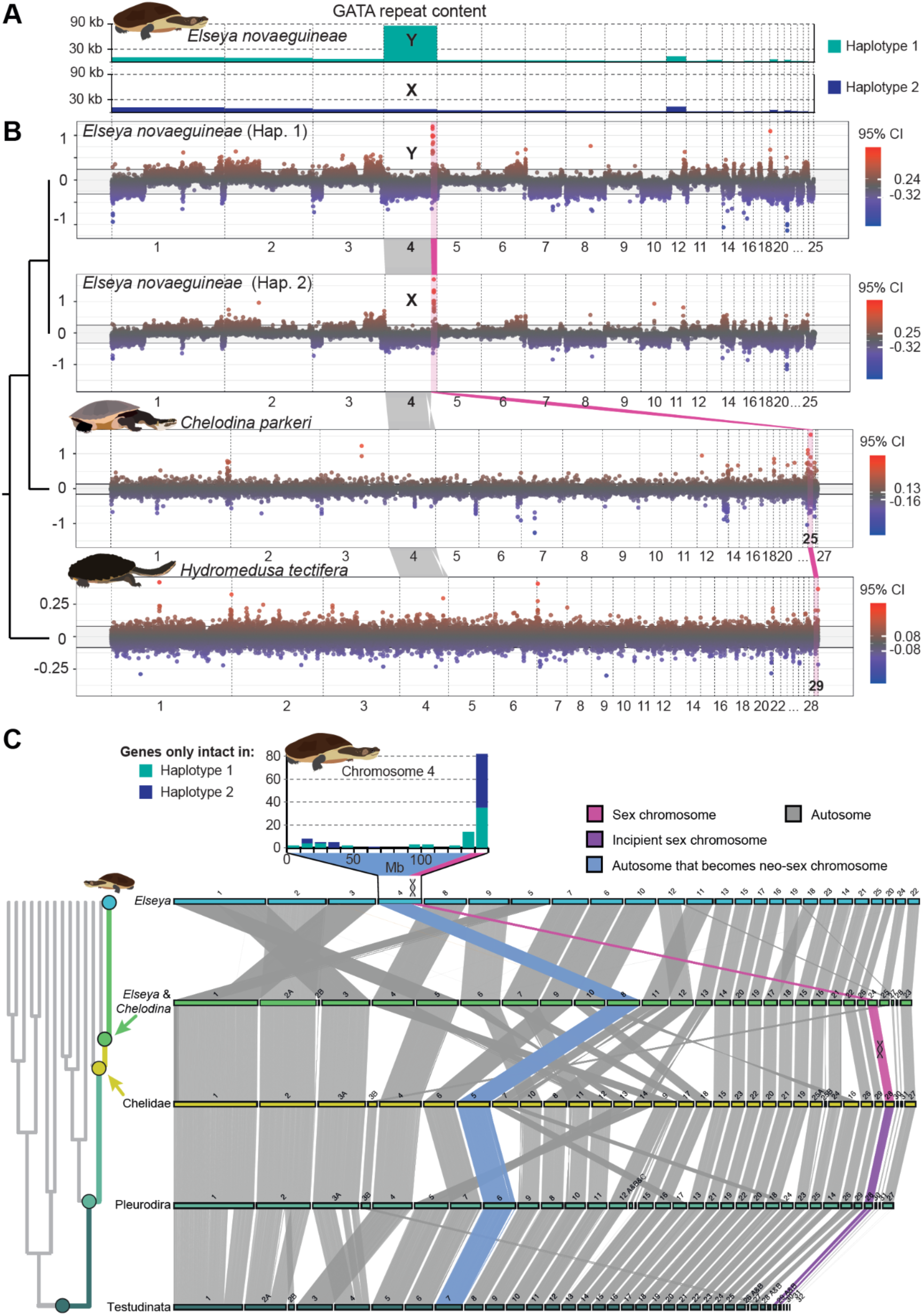
Sex chromosome identification and evolution in Chelidae. A) Total GATA repeat content in kb, across the chromosome-size scaffolds in both haplotypes of the genome assembly of *Elseya novaeguineae*. B) Genome−wide sex differences in heterozygosity (heterogametic−homogametic) in 50 kb windows for both haplotypes of *Elseya novaeguineae*, *Chelodina parkeri* and *Hydromedusa tectifera*. Color indicates outliers based on 95% confidence intervals. Chromosomes are named and ordered by size, except for *Elseya* haplotype 1, which was reordered, so that orthologous genomic regions of both *Elseya* haplotypes are illustrated in the same order and orientation. Pink ribbons highlight and connect genomic regions orthologous to the outlier region in *Elseya novaeguineae* across all species. Grey ribbons connect genomic regions orthologous to the rest of chromosome 4 in *Elseya* across all species. C) Reconstructed chromosome evolution from the last common ancestor of all turtles to *Elseya*. Evolution of chromosomes that will become or fuse to sex chromosomes in Chelidae are highlighted in color, while evolution of the remaining autosomes is shown in grey. The large neo-sex chromosome in *Elseya* evolved by a fusion of the ancestral micro-sex chromosome in Chelidae to a large autosome. For chromosome 4 of *Elseya*, a bar plot shows the position (in 10 Mb windows) of genes classified as intact in only one of the two haplotypes.

**Figure 4:**
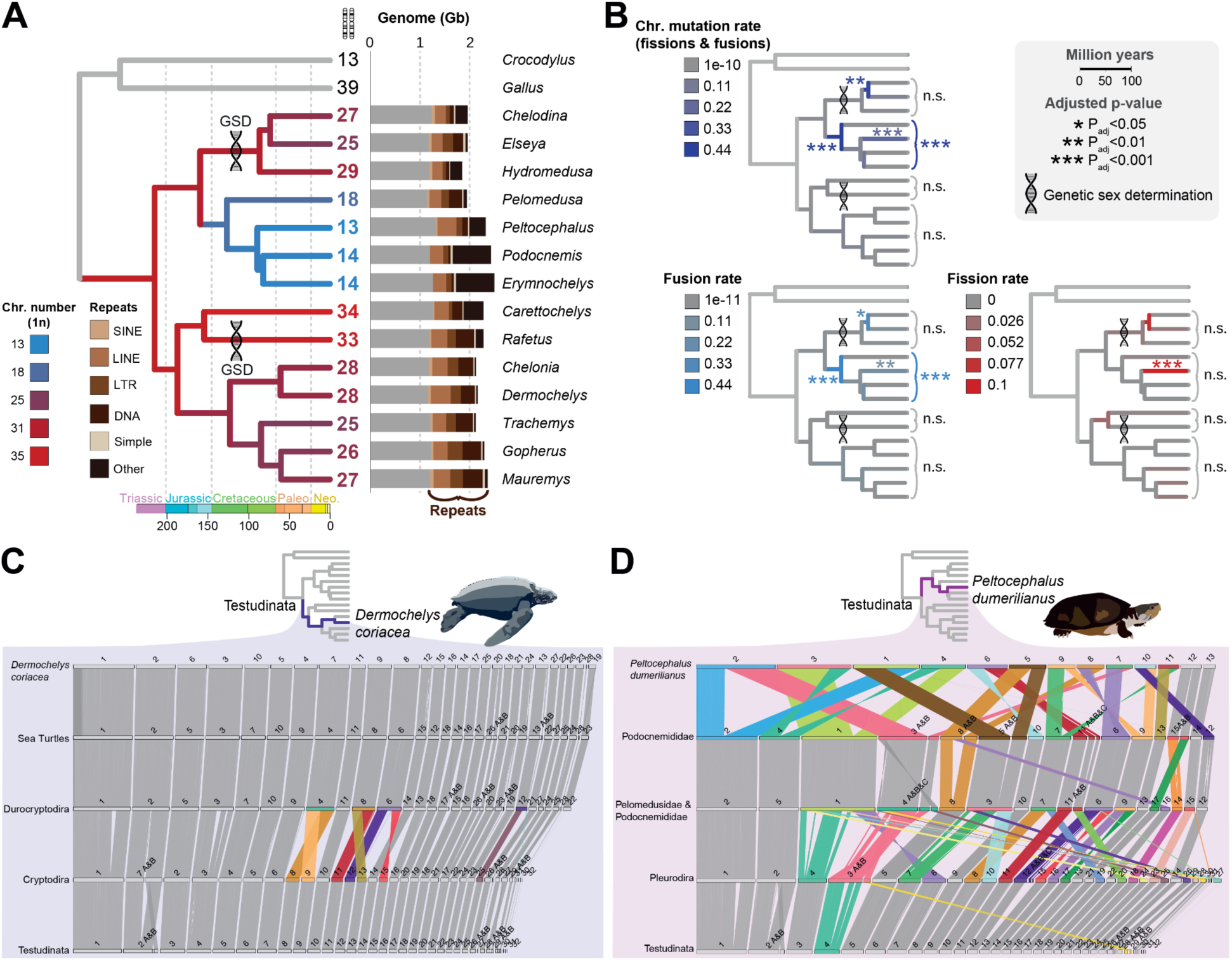
Chromosome evolution in turtles. A) Chromosome number evolution across turtles. The color of phylogenetic branches and numbers at the tips illustrate chromosome numbers and double helixes show when genetic sex determination (GSD) evolved. Bar plots show genome size and repeat content (brown) for each species. B) Chromosome mutation rates (purple, top), fusion rates (blue, bottom left) and fission rates (red, bottom right) across turtles. Asterisks indicate on which branches and in which clades these rates are significantly accelerated. C, D) The trajectory of chromosome evolution from the reconstructed last common ancestor of extant turtles to two representative species, the cryptodiran *D. coriacea* (C) and the pleurodiran *P. dumerilianus* (D). Breaking and fusing chromosomes are highlighted in color.

Second, since the heterozygosity outlier region in *Elseya novaeguineae* is found in both haplotypes of the haplotype-resolved male assembly, we analyzed the abundance of GATA DNA sequence motifs to identify which haplotype contains the X/Y chromosome. This analysis is based on the reported observation that GATA motifs accumulate on the Y chromosome, but not on the X chromosome and the autosomes ^34^. We found a dramatically increased GATA repeat content in chromosome 4 of haplotype 1, strongly suggesting that haplotype 1 contains Y and haplotype 2 contains X chromosome (Figure 4A). Additionally, we found a genome assembly gap situated in a ∼74kb stretch of GATA repeats motifs on the Y chromosome, but not on the X chromosome. Furthermore, there are a number of small unplaced contigs that only consist of GATA repeats, indicating that these contigs originated from an even larger stretch of GATA repeats that could not be fully assembled on the *Elseya* Y chromosome (Figure S2). Third, leveraging the haplotype-resolved *Elseya novaeguineae* assembly, we investigated potential functional divergence of the sex chromosomes. To this end, we used TOGA to identify genes that are intact in one but not the other sex chromosome. This analysis showed a pronounced peak at the end of chromosome 4 (inset in Figure 3C).

Next, we re-sequenced males and females of *Chelodina parkeri* (micro sex chromosome), and *Hydromedusa tectifera* (unknown, likely genetic sex determination system), which represent the other two major Chelid clades and to identify their sex chromosomes (Figure 3A). For *Chelodina*, the most pronounced increase in heterozygosity in males compared to females was found on a microchromosome: chromosome 25 (Figure 4B). This is in line with previously proposed homomorphic X/Y micro-sex chromosomes in *Chelodina* ^32,34^. For *Hydromedusa*, having unknown sex chromosomes, we found less pronounced outlier regions compared to the other two species. The three most extreme outlier-windows were situated on chromosome 1, chromosome 7, and chromosome 29.

In combination, we identify and put into genomic context both micro and macro sex chromosomes in Chelidae including both the X and the Y chromosome of *Elseya* and thereby confirm the sex chromosome diversity in Chelidae.

### Genetic sex determination in Chelidae evolved once on a micro sex chromosome

Different sex chromosomes in Chelidae raise the question whether GSD evolved multiple times in Chelidae. To illuminate the origin of GSD and the evolution of sex chromosomes in Chelidae, we reconstructed and traced the evolution of ancestral chromosomes from the last common ancestor of all turtles to the 3 chelid species *Elseya novaeguineae, Chelodina parkeri* and *Hydromedusa tectifera*.

To investigate whether the outlier windows on different chromosomes represent distinct or homologous genomic regions, we generated whole genome alignments, which revealed that the chromosome 4 *Elseya novaeguineae* outlier region is homologous to chromosome 25 of *Chelodina parkeri* (the clearest sex chromosome candidate) and chromosome 29 of *Hydromedusa tectifera* (one of three candidates). This result indicates a single origin of their sex chromosomes, followed by genomic rearrangements. To confirm this, we mapped the male/female read data from *Chelodina* and *Hydromedusa* to the *Elseya* reference genome. In both cases, the highest outlier window was located in the outlier region on chromosome 4 of *Elseya* (Figure S3). Thus, our results provide strong support for micro-sex chromosomes in *Chelodina* and *Hydromedusa* and indicate that one homologous region contains sex linked-regions across all three species. Interestingly, in *Hydromedusa tectifera* we also observed a second outlier region at the start of chromosome 7, which calls for future investigations to explore whether this reflects a more recent second sex chromosome or a turnover event (Figure 3, S3).

To investigate the ancestry of the Chelidae sex chromosome, we employed ancestral genome reconstructions with AGORA ^68^ to infer chromosome evolution since the last common ancestor of all living turtles. Our results indicated that the micro sex chromosome is ancestral and that the macro-sex chromosome of *Elseya* evolved through a fusion of this ancestral micro sex chromosome to a large autosome (Figure 3C). In contrast to previous hypotheses ^35^, we found that the larger size of the Y compared to the X chromosome in *Elseya* and its sister genus *Emydura* is not caused by a loss of the ancestral micro-X chromosome (Figure S4). Instead, both the ancestral micro-Y and micro-X fused to a large pair of autosomes. More specifically, our analyses reveal that in the lineage leading to *Elseya* the ancestral pair of micro sex chromosomes fused to the autosomal chromosome pair 8 (Figure 3C). Combined with our analyses of GATA repeats on the sex chromosomes, this indicates that instead of an ancestral micro-X chromosome loss, the larger size of the Y chromosome compared to the X chromosome is mainly driven by repeat content increase on the Y.

These findings resolve the long-standing debate about the origin of GSD and sex chromosome diversity in Chelidae by providing strong evidence that GSD evolved once on a micro-sex chromosome in Chelidae more than 80 million years ago which later fused to a macro-autosome in the ancestor of *Emydura* and *Elseya*.

### Chromosome evolution is punctuated in turtles

To explore how the remarkable karyotype diversity (ranging from n=13 to n=34) in turtles evolved, we used our new Pleurodira reference genomes and previously published high-quality genome assemblies ^43,45,58,59,69^ to reconstruct turtle chromosome evolution from the last common ancestor of all living turtles to 14 extant turtles using AGORA ^68^. The included species cover all major turtle clades and the full range of karyotype diversity across turtles.

Our ancestral reconstructions contain a minimum of 11,925 conserved one-to-one orthologous genes in curated chromosomes and reveal a relatively high chromosome number in the last common ancestor of extant turtles (n = 32) with subsequent reduction in chromosome numbers in most clades. This result based on reconstructed chromosome evolution should be more reliable than previous inferences based on maximum likelihood reconstruction indicating only n = 26 chromosomes in the last common ancestor of turtles^70^. Furthermore, excluding the possibility that chromosome arms were falsely reconstructed as separate chromosomes we find that each of the inferred turtle LCA chromosomes are conserved in at least one species in our dataset (Table S2).

Estimating the rate of events per million years, we found generally low fission (∼0.01 fissions/My) and fusion (∼0.03 fissions/My) rates for almost all branches in the turtle phylogeny, indicating comparably slow genome evolution, which is similar to frogs ^71^. However, our analysis also discovered punctuated chromosome evolution in turtles with up to ∼10-fold increased fission and up to ∼15-fold increased fusion rates (Figure 4B). The most extreme change in chromosome numbers is a reduction from n=31 to n=17 chromosomes in ∼33 My leading to the common ancestor of Podocnemididae and Pelomeduside. Accordingly, different evolutionary lineages exhibit pronounced differences in chromosome evolution rates (Figure 4C, D). While the 32 chromosomes of the last common ancestor of all turtles remained largely conserved for >200 My in the leatherback (4 fusions) or the softshell turtles (1-2 fissions), within the same time period chromosomes of *P. dumerilianus* underwent 26 fusions and 9 fissions. Throughout time, significant accelerations of chromosome evolution only occurred on some branches and were not linked to any obvious major bioclimatic events (Figure 4 A,B).

Although fissions were generally rare in our dataset (n=17), we observed a recurrent fission point on stem-turtle chromosome five, which broke in three lineages independently (Tables S2, S3). Recurrent breakpoints have previously been found in plants ^72^ and animals ^73–75^. Additionally, more than half of all fissions occurred in the terminal branch leading to *Peltocephalus dumerilianus*, with four fissions occurring precisely at previous chromosome fusion points. Such sites may be particularly prone to fission because recent fusion junctions can remain structurally fragile and because purifying selection is likely weaker at these than at other genomic regions where fissions would disrupt long-established regulatory interactions. Furthermore, our results underline the importance of reconstructing chromosome evolution instead of inferring the number of fission and fusion events based on phylogenetic inference of ancestral chromosome numbers. For example, even if ancestral karyotypes were correctly inferred, a reduction of 15 to 13 chromosomes in the branch from Podocnemididae to *Peltocephalus dumerilianus* would only be associated with two fusions. However, this would represent a 10-fold underestimation of the actual number of events comprising 11 fusions and 9 fissions (Figure 4D). Consistent with recent findings of punctuated chromosome evolution in other clades ^71,74–76^ and slow molecular evolution of turtles ^77,78^, our analyses indicate that chromosome evolution is slow in turtles, but interrupted by periods of massive rearrangements.

### Sex determination and chromosome number do not coevolve in turtles

Previous studies reported that chromosome number and sex determination coevolve in turtles. This finding was based on phylogenetic reconstruction of sex determination modes and inferred ancestral chromosome numbers ^26,66^ and primarily driven by higher chromosome numbers in Chelidae (GSD) and softshell turtles (GSD, but TSD in *Carettochelys*), compared to their TSD sister clades. Under this hypothesis, either selection favoring sex determination turnover also promotes genomic rearrangements, or chromosome evolution facilitates transitions in sex-determining mechanisms by altering gene regulation ^66^. To test whether chromosome number and sex determination coevolve, we build on our finding that GSD likely evolved once in stem-Chelidae and used our ancestral chromosome reconstructions covering two independent cases of GSD evolution to estimate whether chromosome evolution rates or karyotypes changed significantly following the evolution of GSD.

In contrast to previous hypotheses, we found no evidence that chromosome number and sex determination coevolve in turtles. Although chromosome evolution is accelerated on several branches (Figure 4B), neither changes in karyotype, nor chromosome evolution rates measured as fission, fusion or combined fission+fusion rates were accelerated in the phylogenetic branches in which GSD evolved or across GSD clades. Instead, the previously reported higher chromosome numbers in GSD turtles appears to be primarily due to a reduction in chromosome numbers in the sister clade of Chelidae, which includes the families Pelomedusidae and Podocnemididae, where only TSD has been reported.

Because fusions between sex chromosomes and autosomes can resolve intralocus sexual antagonism and were shown to be accelerated in some animals ^79^, we estimated the likelihood of the observed sex chromosome autosome fusion in Chelidae. This analysis revealed that observing the single sex chromosome autosome fusion in Chelidae is not unexpected (>50% chance), given the karyotype, the sex determination system and the total number of fusion events. Thus, we cannot find any support for the hypothesis that sex determination and chromosome number coevolve in turtles.

### Evidence for repeat content promoting chromosome fissions

Our analyses indicated that chromosome evolution is not explained by sex determination modes in turtles, which raises the question what drives lineage-specific chromosome evolution rates. Genomic regions that are prone to breaking have previously been linked to open chromatin, high gene density and increased repeat content ^80,81^. However, it is difficult to distinguish between causal factors that drive chromosome fissions and correlated factors that may occur as a consequence of fissions. For example, lineage-specific increases in repeat content may directly promote and thus precede chromosome fission events by serving as templates for non-homologous allelic recombination, or occur as a consequence of chromosome fission events due to relaxed selection and the addition of telomeric repeats ^80,82,83^.

We used the only turtle lineage with significantly increased fission rates, *Peltocephalus dumerilianus*, as a test case to explore whether increased repeat content is a potential contributing factor (Figure 4B, D). Our premise is as follows. If repeat content promoted fissions in *Peltocephalus*, its breakpoint regions should exhibit increased repeat content not only compared to other genomic regions, but also compared to other taxa. Furthermore, if repeat content promotes fissions, local repeat content increases should precede fission events, which can be detected as repeat content increases in the *Peltocephalus* sister clade. In contrast, if increased repeat content near the breakpoints is a consequence of fissions, then it should occur after the fission events and be limited to *Peltocephalus*. Here we used the sister lineage of *Peltocephalus* where the chromosome never broke to distinguish between the two scenarios.

We analyzed the repeat content in breakpoint regions and the genomic background in a phylogenetic context to explore 5 non-mutually-exclusive hypotheses that should result in different patterns of repeat contents across the genome and across the phylogeny as illustrated in Figure 5A: i) no correlation exists between breakpoints and repeat content, ii) repeat content is generally increased in breakpoint regions, but not linked to accelerated fission rates, iii) increased repeat breakpoint content is a consequence of chromosome fissions, iv) local repeat content increases the chance for chromosome fissions, and v) global repeat content increases the chance for chromosome fissions.

**Figure 5:**
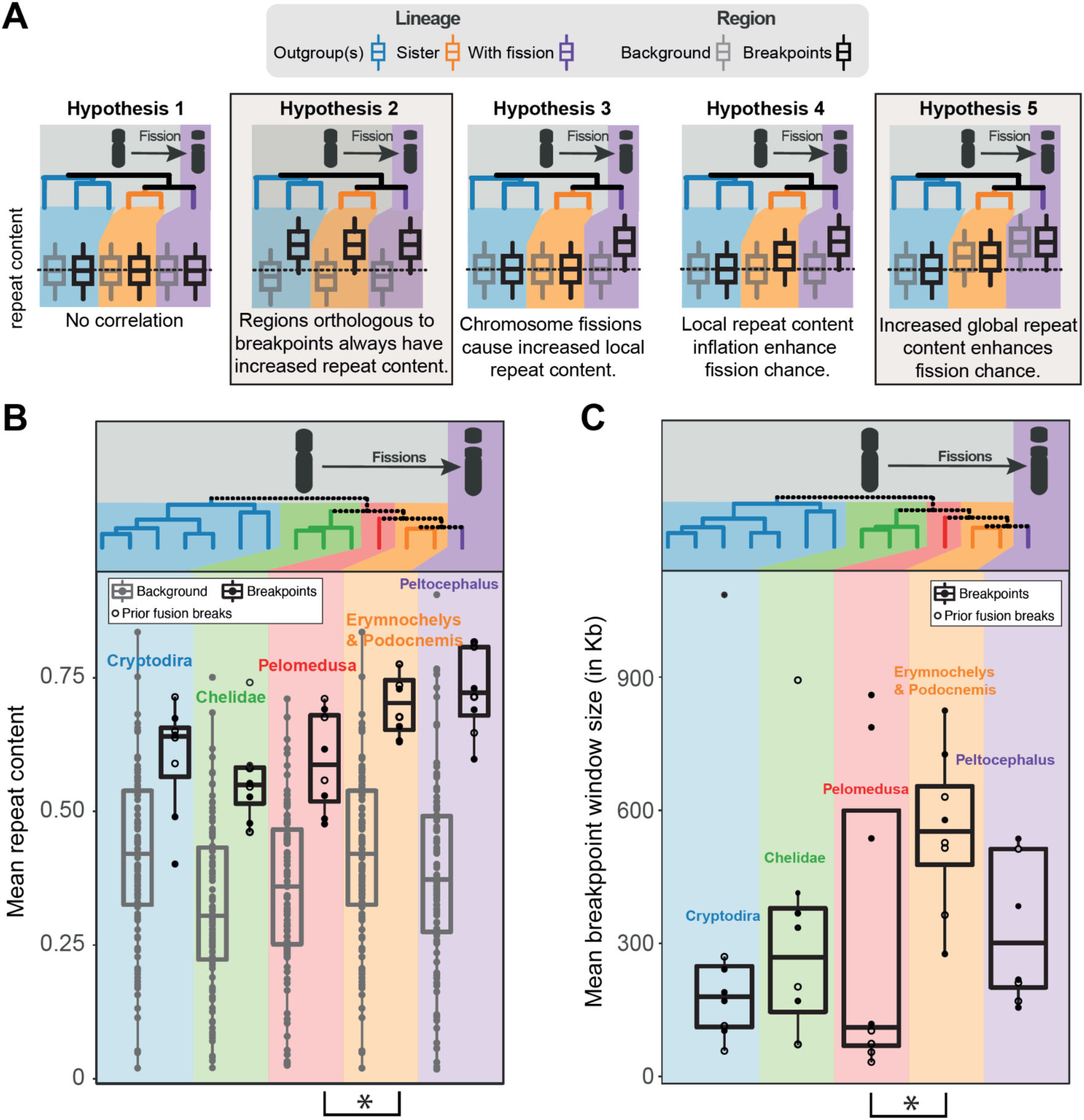
Chromosome breakpoint repeat content evolution in turtles. For five different hypotheses A) illustrates the expected repeat contents across regions (breakpoints vs genomic background) and phylogenetic clades. Boxplots in B) show the repeat content for eight chromosome breakpoints (black) identified in *Peltocephalus* (purple) and 100 genomic background windows (grey) across five turtle clades. For clades with multiple species the mean repeat content is shown for each window. Breakpoints that are previous fusion points are shown as empty circles. For the same clades and breakpoints, C) shows the breakpoint window sizes.

In line with hypothesis ii) and previous work in other organisms ^80,81^, we found repeat content to be generally increased at breakpoint regions compared to the genomic background across all turtles (Figure 5B). Although *Peltocephalus*, in which the fissions occurred, had the highest breakpoint repeat content, it was not significantly increased (p-value = 0.3115) compared to its sister lineage (*Erymnochelys* and *Podocnemis*). Notably, in this sister lineage, breakpoint repeat content was significantly increased compared to the outgroup (*Pelomedusa*: p-value = 0.035) (Figure 5B). Additionally, significantly increased breakpoint region sizes in the sister clade compared to the outgroup (*Pelomedusa*: p-value = 0.038) support increased repeat content already inflated the breakpoint regions before *Peltocephalus* split from its sister clade (*Erymnochelys* and *Podocnemis*). In line with hypothesis v), this increased repeat content is not locally restricted to breakpoints, but roughly proportional to increasing genome-wide repeat contents. While the breakpoint regions do gain more repeats than other parts of the genome, the proportions of repeat contents inside and outside of breakpoint regions remain roughly the same (repeat content breakpoints/background: 1.67 in *Pelomedusa* (outgroup), 1.69 in *Erymnochelys* and *Podocnemis* (sister), 1.89 in *Podocnemis* (fissioned)) (Figure S5).

Our observation of recurrent breakpoints and locally elevated repeat content add further support to the hypothesis that chromosome breaks do not occur randomly across the genome but accumulate at “fragile” genomic regions. These results support a scenario where a genome-wide repeat content increase promoted chromosome fissions by increasing the repeat content in already repeat-rich fragile genomic regions.

### Gene losses are associated with innovation and adaptation in turtles

Gene losses are an often-underappreciated source of variation that contributed to several evolutionary adaptations ^84–88^. To explore the potential contribution of gene losses to evolutionary innovations in turtles, we used TOGA ^47^ to identify genes lost in all extant turtle genomes and inferred those that were likely lost in the stem turtle lineage.

In total, we identified 12 genes that were lost in stem turtles, but intact in chicken and crocodiles. Intriguingly, defects in two of these 12 genes (*PRKG2, MATN3*) are known to impair skeletal development in mammals by causing disproportionate dwarfism with short, broad body shapes ^89–96^. Only a total of 72 genes are known to cause disproportionate dwarfism, hence observing two such genes among the 12 turtle-lost genes by chance is highly unlikely and corresponds to a ∼54-fold enrichment (p < 0.001, hypergeometric test, “disproportionate dwarf”) ^97,98^.

The first gene, *PRKG2*, encodes for a cGMP-dependent protein kinase that plays a key role for chondrogenic differentiation and endochondral bone growth. Its loss coincides with a genomic deletion in stem turtles comprising the last seven exons (Figure S6). Loss of function mutations in *PRKG2* cause disproportionate dwarfism with short, broad body shapes across a wide range of mammals including mouse, rat, dog, cow and human ^89–93^. For example, *PRKG2* knockout in rats causes significant rostroventral shortening with shortened skull, femur, vertebra and limb bones, while skull and clavicle width remain constant.

The second turtle-lost disproportionate dwarfism-associated gene, *MATN3*, encodes a filament-forming extracellular matrix protein that is primarily expressed in cartilage and bone ^102^. In humans, mutations in *MATN3* cause skeletal disorders such as multiple epiphyseal dysplasia and spondyloepimetaphyseal dysplasia, which are characterized by impaired long bone growth leading to disproportionate dwarfism with short limbs and bowed legs ^94–96,103^. It appears plausible that loss of *PRKG2* and *MATN3* contributed to the evolution of a rostrocaudally compressed skeleton in stem-turtles.

Furthermore, *MATN3* loss may be involved in turtle-specific differences in rib ossification. In the extracellular matrix, MATN3 acts as an antagonist of BMP2 ^104^, a secreted morphogen ^104^ hat is critical for shell patterning and endochondral rib ossification ^105,106^. Importantly, the bony plate at the base of turtle carapace (the costal plate) develops from the ribs via an expanding periosteum that leads to an exaggerated outward rib-growth ^101^, which may be combined with BMP-induced dermal ossification ^107,108^. *BMP2* can induce the formation of periosteum-like tissues ^109^ and overexpression of *BMP2* results in ectopic bone formation close to ribs, scapulae, and neck in mice ^110^. Thus, removing the suppressive effect of *MATN3* on *BMP2* signaling during rib ossification may promote *BMP2*-induced expansion and ossification of the periosteum and potentially dermal ossification (Figure 6E).

**Figure 6:**
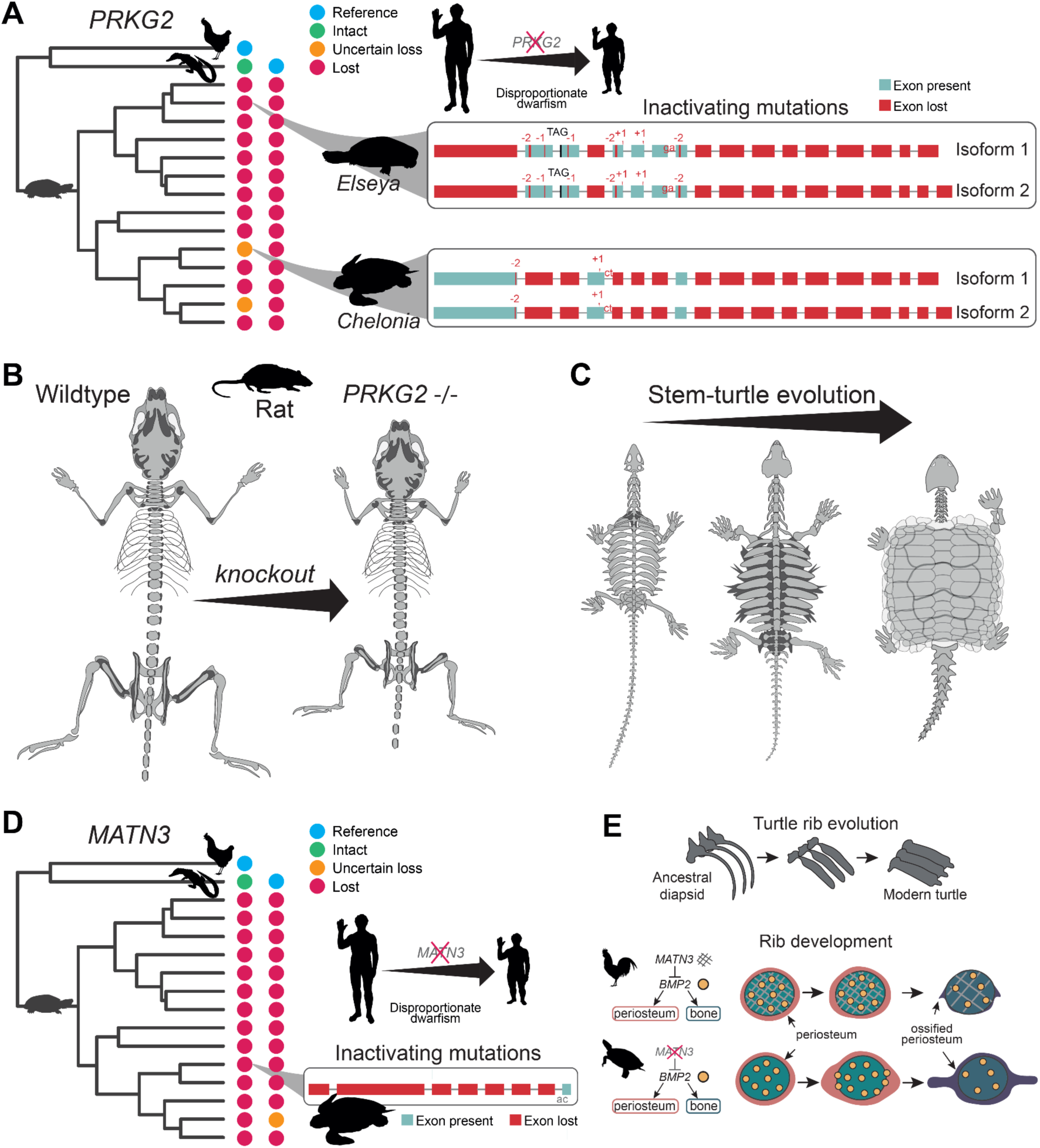
Gene losses associated with skeletal transformation in turtles. A) *PRKG2* is lost in turtles. Using chicken and crocodile as references, colored circles tips show the gene loss pattern in 14 high-quality turtle genomes across the turtle phylogeny using chicken (left column) and crocodile (right column) as references. Inactivating mutations are shown in both isoforms for one representative side-necked and hidden neck turtle. *PRKG2* loss causes disproportionate dwarfism in humans and many other mammals. B,C) PRKG2 knockout caused rostrocaudal compression of the skeleton in rats (redrawn based on ^99^), which resembles skeletal evolution in stem turtles shown in C) (redrawn based on ^100^). D) Similar to *PRKG2* loss shown above, *MATN3* is lost in turtles, which also causes disproportionate dwarfism in humans. E) Turtle rib evolution is shown together with a hypothesis how *MATN3* loss may contribute to turtle specific ossification during rib development. Here, rib development is shown for chicken and turtle and we hypothesize that removing the suppressive effect of *MATN3* on *BMP2* signaling may promote turtle-specific periosteum formation and later ossification as previously observed ^101^.

Beyond skeletal development, we also detected a loss of *ADGRF2* (also known as *GPR111*), a gene required for proper enamel formation. *ADGRF2* modulates expression of the enamel forming *KLK4* ^111^, which has been found to be lost in turtles, similar to other genes involved in tooth formation such as *ENAM, AMEL*, *AMBN*, and *DSPP* ^15,59^. Therefore, *ADGRF2* loss is likely linked to teeth loss in the ancestor of all extant turtles.

Additionally, turtles have lost two genes, *EPB42* and *PBLD,* with important roles for blood vessel formation and oxygen transport. *EPB42* has erythrocyte-specific expression (Figure S7, https://www.proteinatlas.org/) and impaired human *EPB42* function causes spherical shape in red blood cells (spherocytosis) ^112^, which resembles features of turtle erythrocytes that are exceptionally large and oval ^113,114^. *PBLD* inhibits hypoxia-induced angiogenesis, and loss of *PBLD* may help to promote angiogenesis in heart and brain to maintain blood flow to these vital organs during anoxia ^25,115^.

Notably, our data further revealed that turtles lost the redox sensor *NMRAL1* (also known as *HSCARG*), which plays an important role for DNA damage response and cell cycle control ^116–118^ (Figure 7A). In response to oxidative stress, NMRAL1 dimers disaggregate into monomers, which enter the nucleus and downregulate translesion synthesis, a DNA damage tolerance mechanism ^116^ (Figure 7B). *NMRAL1* knockout traps repair proteins at lesion points and continuously activates checkpoint signaling, resulting in reduced mutation rates and prolonged cell cycle duration, which limits the oncogenic transformation capacity of cells having cancer-causing mutations ^117,119^. Consistently, overexpression of *NMRAL1* exacerbates mammary tumorigenesis ^116^. Interestingly, several studies reported slow cell cycles ^120,121^ and increased cancer resistance in turtles ^11,12,17^, which correlates with *NMRAL1* loss, and the loss of several other genes (*DHX32*, *PBLD*, *GPR107*, *BDKRB1*) known to affect cancer progression ^122–127^. *NMRAL1* links oxidative stress to a reduced DNA damage response ^118^, which may have been detrimental for stem turtles which were exposed to increased levels of oxidative stress. Thus, it appears plausible that inactivation of *NMRAL1* helped to reduce mutation rates in stem-turtles and contributed to slower cell cycles and increased cancer resistance.

**Figure 7:**
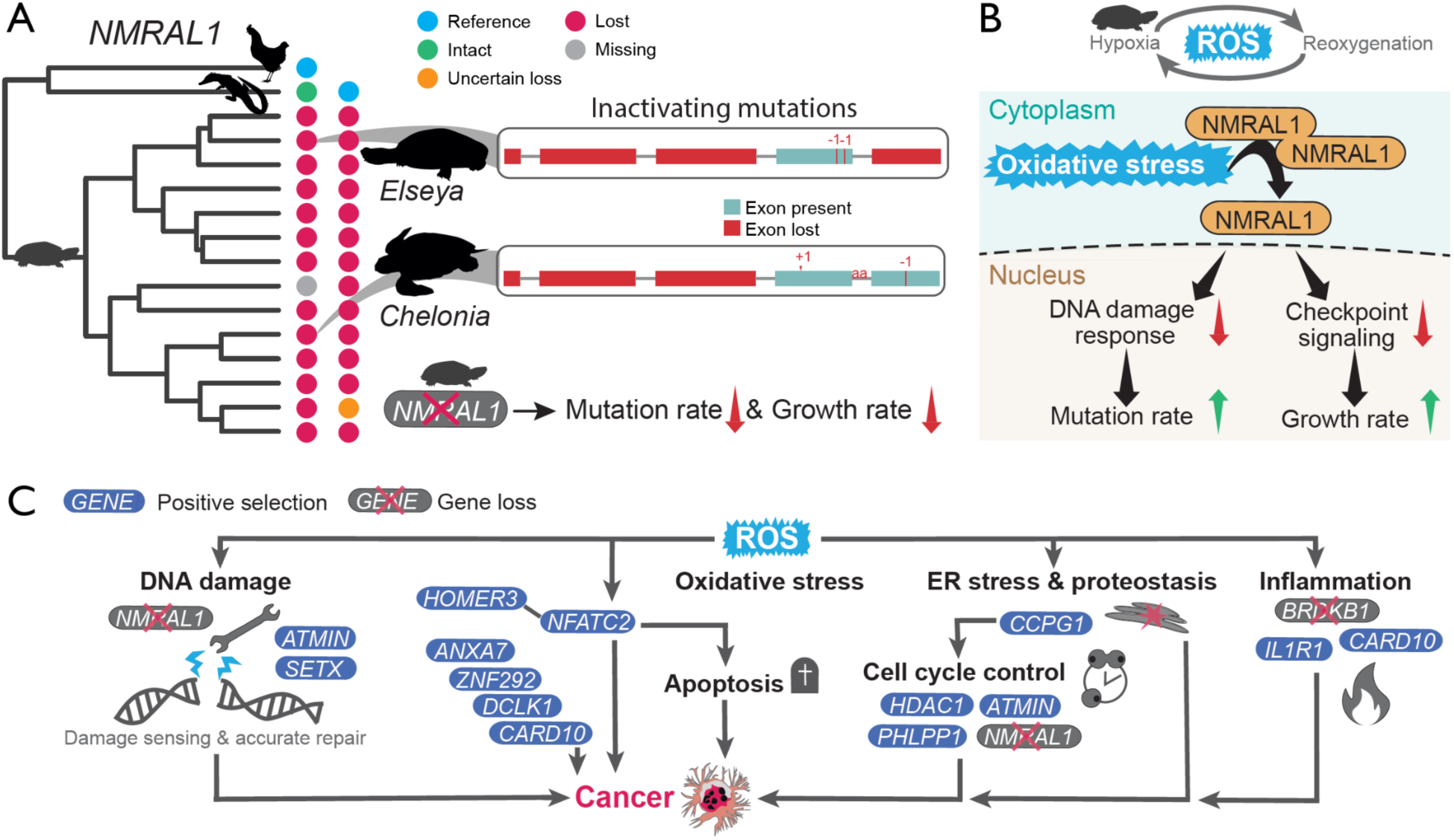
*Adaptive evolution* associated with cancer resistance under oxidative stress in turtles. A) *NMRAL1* is lost in turtles. Using chicken (left column) and crocodile (right column) as references, colored circles show the gene loss pattern in 14 high-quality turtle genomes across the turtle phylogeny. Inactivating mutations are shown for one side-necked turtle and one hidden neck turtle. *NMRAL1*-loss causes decreased mutation rates and decreased growth rates. B) *NMRAL1* signaling typically downregulates the DNA damage response and decreases checkpoint signaling in response to oxidative stress. C) Gene losses and genes under positive selection in the stem turtle lineage are associated with key pathways linking oxidative stress to cancer occurrence and progression.

To ensure that our findings are not biased towards frequently lost genes, we explored whether the 12 genes lost in turtles are conserved in other tetrapods. Analyzing high-quality genomes of 651 mammals and 107 birds revealed that the turtle-lost genes are highly conserved in birds and mammals (Figures S8-S12). The only exception is *NMRAL1*, which is highly conserved in mammals, but more frequently inactivated in birds (Figure S10). This could be related to increased levels of oxidative stress due to flight ^128^. Interestingly, the bird clades that seem to have lost *NRMAL1* typically have long lifespans such as the exceptionally long-lived parrots (Figure S10)^129^. In combination, our results reveal clear associations of losses of otherwise conserved genes with both morphological and physiological adaptation that shaped the evolution of turtles.

### Adaptation of DNA repair, stress signaling and cell-cycle control likely promotes turtle cancer resistance

Prompted by our finding that gene losses may have contributed to enhanced cancer resistance in turtles, we next investigated genes evolving under positive selection. A systematic screen identified 155 genes with significant signatures of positive selection in the stem-turtle lineage (Table S4). Several of these genes function in DNA damage response, oxidative stress signaling, proteostasis, and tumorigenesis; processes particularly relevant for long-lived, cancer-resistant turtles that are exposed to repeated hypoxia–reoxygenation cycles (Figure 7C).

Among DNA damage response factors, we detected positive selection in *ATMIN*, an oxidative stress–responsive transcription factor required for activation of the ATM kinase dependent repair of oxidative stress induced DNA damage ^130,131^, and in *SETX*, a BRCA1-recruited helicase that resolves co-transcriptional RNA:DNA R-loops to prevent double-strand breaks and replication fork stalling ^132,133^. Adaptive changes in these genes may enhance lesion detection and genome maintenance under oxidative stress.

Chronic oxidative stress promotes protein misfolding ^134^. We detected positive selection in *CCPG1*, an ER-phagy receptor induced by the unfolded protein response that maintains proteostasis by targeting stressed endoplasmic reticulum regions for lysosomal degradation ^135–137^. Altered CCPG1 function could help to mitigate ER stress and initiate ER stress–induced cell-cycle arrest ^136^, thus limiting the proliferation of persistently damaged cells.

Recurrent oxidative stress can further induce chronic inflammation, a hallmark of aging and tumorigenesis that promotes every stage of tumor formation ^138,139^. In addition to the loss of the stress-induced bradykinin receptor *BRDKB1*, which activates pro-inflammatory signaling via several cytokines such as IL-1β ^140^, we discovered signatures of positive selection on the IL-1β receptor *IL1R1,* a mediator of IL-1β driven inflammatory signaling during tumor formation and progression ^141^ and *CARD10,* which drives tumor growth and invasiveness by activating NF-κB signaling ^142–144^. Adaptive changes in these pathways may mitigate chronic inflammation driving tumorigenesis and cancer progression.

Several cancer-related genes also exhibit signatures of adaptive evolution. These include *NFATC2*, encoding an oxidative stress–activated transcription factor implicated in mitochondrial dysfunction, inflammation, and apoptosis, and its regulator *HOMER3*, which both promote lung tumor growth and metastasis ^145–149^. In addition, we observe positive selection in *PHLPP1*, encoding a phosphatase that suppresses the tumor-promoting ATK kinase ^150^; *HDAC1*, encoding a histone deacetylase that promotes degradation of the p53 tumor suppressor and represses cell-cycle gene transcription in an Rb-dependent manner ^151,152^; *ZNF292*, a zinc finger transcription factor that restricts cell proliferation by p27 stabilization ^153^; *ANXA7*, a calcium-activated GTPase with tumor suppressor activity ^154,155^; and *DCLK1*, a hypoxia-induced kinase regulating cancer stemness, epithelial–mesenchymal transition, and malignancy ^156^. Together, these patterns suggest adaptive modulation of pathways involved in tumor suppression, DNA repair, cellular proliferation control and stress tolerance in the turtle lineage.

## Discussion

To overcome the previous lack of reference-quality Pleurodira genomes and illuminate turtle genome evolution, we provide seven highly complete chromosome-level genomes. With these assemblies, we substantially improve the amount and quality of available genomic resources for Pleurodira and turtles as a whole and facilitate future work on turtle conservation genomics and genome evolution.

Using our new resources, we generated a highly supported, well resolved turtle phylogeny based on the largest set of molecular markers, which supports recent phylogenetic hypotheses including the previously debated sister relationship of *Platysternon* and Emydidae ^42,49^.

For the first time, we identify the sex chromosomes of *Hydromedusa* and *Chelodina*, anchor the sex chromosome of *Elseya* in genomic context and use a combination of genome alignments and ancestral chromosome reconstructions to resolve the evolution of GSD in Chelidae: GSD evolved only once in Chelidae >80 million years ago on a micro-sex chromosome and both the micro-Y and the micro-X later fused to a macro-autosome resulting in a macro neo-sex chromosome in *Emydura* and *Elseya*. Despite their age (>80 My), chelid (micro)-sex chromosomes remained highly homomorphic, which adds to findings of old, homomorphic sex chromosomes in other species like ratite birds and boas ^157,158^, and challenges the traditional view that Y and W chromosomes are born to die ^159,160^.

Beyond sex chromosomes, we discovered that punctuated chromosome evolution with rare, but pronounced (up to 15-fold) accelerations of otherwise slow fission and fusion rates gave rise to the remarkable karyotype diversity in turtles. While similar patterns were recently discovered in butterflies ^76^, denser chromosome-level genome coverage in other taxonomic groups will enable systematic studies to clarify whether short and pronounced bursts are rare events or a general evolutionary mode of chromosome evolution. Additionally, illuminating what underlies such bursts of chromosome evolution is fundamental to understanding the drivers of chromosome evolution ^80^. We refute the long-standing hypothesis that accelerated chromosome evolution is linked to transitions between sex determination modes in turtles ^26,66^. In contrast, we find patterns consistent with repeat content promoting chromosome fission rates. While increased repeat content is a common feature of breakpoint regions ^75,80,81^, it has been debated whether repeats contribute to fragility or whether they are primarily a byproduct of other, more decisive features such as 3D conformation and open chromatin ^81^. Our results are in line with elevated genome-wide repeat content contributing to increased fission rates by further increasing local repeat contents at already repeat rich fragile genomic regions, which can increase the chance for non-homologous allelic recombination ^80,82^. Comparative analyses across a wider taxonomic range of organisms comprising varying evolutionary rates will uncover whether bursts of chromosomal evolution typically coincide with elevated repeat content and, if so, why some species with extremely repeat rich genomes retain conserved karyotypes ^71,161^.

Today, turtles are one of the most endangered vertebrate clades. To better understand what drives recent turtle declines and illuminate how turtles responded to historical climate change, we inferred population histories of side necked turtles. Our results show clear effects of historical climate change in the distant past, which differed between continents.

For the most recent time since the last interglacial, PSMC consistently inferred dramatic population declines for all of our species. As climate is discussed as one factor contributing to future turtle decline ^37,42^, one might expect that inferred declines would be related to climate fluctuations as well. However, similar declines can be inferred solely because of existing population structure ^64^. To differentiate between these scenarios, we use GLMMs to statistically test whether expected drivers of population declines explain variation beyond the gradual decline generated by population structure. Notably, time and thus population structure alone was sufficient to explain the observed population declines. While this does not rule out an effect of climate, biome, or type of sex-determination, population structure likely dominates declining population size estimations in the recent past, which makes it impossible to detect potential additional effects. This result is important, because similar declines in the inverse instantaneous coalescence rate are extremely common and are often interpreted as biologically meaningful (e.g., climate-/human-induced) changes in population size ^162,163^. If climate change alone does not explain well the decline of organisms, disregarding the potential effect of population structure, may lead researchers to erroneously attribute population declines to additional factors, e.g., human hunting pressure ^162^. Thus, our results add to theoretical work ^64^ and call for more careful interpretation of putative drivers of relatively recent (here ∼100,000 years) population declines.

Finally, to gain insight into the molecular evolution associated with turtle innovations, we investigated an often-overlooked molecular mechanism: gene losses. Interestingly, our results reveal several gene losses associated with evolutionary innovation in turtles, including their highly derived body plan with a rostrocaudally compressed skeleton (*PRKG2* and *MATN3*), their toothless beak-like yaw (*ADGRF2*), hypoxia tolerance (*EPB42*, *PBLD*), and cancer resistance under increased oxidative stress (*NMRAL1*). In particular, while *PRKG2* and *MATN3* mutations cause pathological morphological phenotypes in mammals, we hypothesize that the loss of these genes may have contributed to the disproportionate dwarfism-like body-plan and turtle-specific rib ossification patterns in these “hopeful monsters”. Functional tests and investigations of regulatory evolution in stem turtles will help to confirm the effects and evaluate the relative importance of gene losses for turtle evolution.

The functional profiles of lost and positively selected genes suggest that turtle cancer resistance may rely less on eliminating damaged cells and more on limiting cellular damage accumulation, promoting accurate repair and maintaining tightly controlled cell proliferation. While employing different pathways, turtles seem to use a “prevent-and-repair” oriented cancer resistance strategy analogous to that described in naked mole rats ^164–166^, contrasting with apoptosis-centered cancer suppression in elephants ^167^.

Overall, our analyses resolve long-standing questions in turtle genome and sex chromosome evolution, shed light on the molecular basis of cancer resistance and highlight gene loss as a potential driver of evolutionary innovation.

## Materials and Methods

### Samples and permits

We obtained blood samples of captive animals from Turtle Island, Austria, that were continuously kept in Europe since 2010 or before. More specifically, the *Erymnochelys madagascariensis* individual was bred from individuals that were imported from the USA in 2010, the *Peltocephalus dumerilianus* specimen is offspring from individuals imported to Germany in 1997 and later transferred to Turtle Island, Austria, parental animals of *Hydromedusa tectifera* were acquired in 1976 and kept in Austria ever since, and *Chelodina parkeri* was kept at the Stuttgart Zoo in Germany since at least 2001 and transferred to Turtle Island, Austria in 2014. Turtle Island is a category A zoo and holds all necessary permits. The *Podocnemis unifilis* individual was bred at the Zoo Leipzig from animals that hatched in Burgers Zoo Arnheim in 1998 and was transferred to the Zoo Leipzig in 1999. The *Pelomedusa somalica* individual was collected in Somalia, Woqooyi Galbeed, Somaliland (N 9 23 27, E 44 07 12) in 2017 (Permit no. Ref. MOERD/M/I/251/2017). Blood samples from *Elseya novaeguineae*, and additional samples from 5 individuals of *Hydromedusa tectifera* and two individuals of *Chelodina parkeri* for resequencing were obtained from specimens that were kept and bred by private breeders in Czech Republic since before 2010 and carried out under the supervision and with the approval of the Ethics Committee of the Faculty of Science, Charles University, and the Committee for Animal Welfare of the Ministry of Agriculture of the Czech Republic (permit No. MSMT-34426/2019-7). The animal handling was performed by accredited researchers (MR: accreditation CZ03540).

### Tissue sampling and DNA extraction

Nucleated blood was drawn from legs of *Erymnochelys madagascariensis*, *Peltocephalus dumerilianus*, *Pelomedusa somalica, Podocnemis unifilis, Hydromedusa tectifera, Elseya novaeguina*, and *Chelodina parkeri* and immediately snap-frozen in liquid nitrogen (*Erymnochelys* & *Peltocephalus*) or first preserved in 95-100% molecular biology grade Ethanol (circa 50 to 100 ul blood/ml Ethanol), before snap-freezing in liquid nitrogen. All blood samples were stored at -80°C.

We extracted ultra-long high molecular weight genomic DNA (uHMW gDNA) from all samples. For the flash frozen blood samples of *Erymnochelys* and *Peltocephalus*, we followed the Bionano Prep Cell Culture DNA Isolation Protocol (30026 Rev E). In brief, blood cells were diluted in one volume of ice-cold Cell Buffer (derived from the Bionano Prep Blood DNA Isolation kit) and mixed by pipetting. The blood slurry was embedded into agarose plugs and treated with Proteinase K and RNase A. uHMW gDNA was extracted from agarose plugs and purified by drop dialysis against 1x TE. Using a PippinPulse^TM^ (Sage Bioscience), revealed uHMW gDNA with lengths of 100 kb to 500 kb.

For blood preserved in Ethanol (*Pelomedusa, Podocnemis, Hydromedusa, Elseya*, and *Chelodina*) uHMW gDNA was extracted with the Bionano Prep SP Frozen Human Blood DNA Isolation Protocol (30246 Rev C) adapted for nucleated blood preserved in Ethanol. In brief, 100 ul nucleated blood cells were collected by centrifugation (5 min at 2000 rcf) and washed twice in 1x PBS. The washed blood cell pellet was dissolved in an equal volume of cell buffer and lysed by adding Proteinase K. After cell lysis, gDNA in lysis buffer was bound to Nanobind Disks in the presence of Isopropanol. After washing, HMW gDNA was eluted and the integrity of the HMW gDNA determined using the Pippin Pulse^TM^ device (SAGE Bioscience), which showed that the majority of fragments was larger than 50 kb.

For *Elseya novaeguineae*, uHMW gDNA was extracted with the circulomics Nanobind Tissue Big DNA kit (part number NB-900-701-01 v1.0 (11/19)) according to the manufacturer’s instructions. Nucleated blood cells were washed twice in PBS before resuspension in 10 ul of washed blood cells in PBS and lysis with Proteinase K. After RNA removal with RNAse A, gDNA was bound to circulomics Nanobind discs and uHMW gDNA was eluted with elution buffer. DNA integrity was determined by pulse field gel electrophoresis using the Femtopulse (Agilent) revealing fragment lengths of ∼120 Gb.

### PacBio library preparation and genome sequencing

PacBio continuous long read (CLR) libraries were prepared for *Chelodina parkeri, Erymnochelys madagascariensis, Hydromedusa tectifera, Pelomedusa somalica, Peltocephalus dumerilianus,* and *Podocnemis unifilis*. To this end, long insert libraries were prepared according to the guidelines for preparing size-selected >30 kb SMRTbell templates using the SMRTbell express Template kit 2.0. Briefly, ultra-long gDNA was sheared to 75 kb fragments with the MegaRuptor device (Diagenode) and 3-6 µg sheared gDNA was used for library preparation. The PacBio SMRTbell libraries were size-selected for fragments larger than 4,5 kb (*Erymnochelys madagascariensis*, *Peltocephalus dumerilianus*) and 25 kb (*Chelodina parkeri, Hydromedusa tectifera, Pelomedusa somalica, Podocnemis unifilis*) with the BluePippinTM device. All size selected libraries were sequenced for 15 hours on the Pacbio Sequel^®^ II on 8M SMRT™ cells using the Sequel® II Binding 2.0 and the Sequel^®^ II Sequencing 2.0 chemistry.

For *Elseya novaeguineae*, PacBio High Fidelity (HiFi/CCS) libraries were prepared using the HiFi Libraries from Low DNA Input Using SMRTbell Express Template Prep Kit 2.0. Briefly, gDNA was sheared to 8 kb with the MegaRuptor device (Diagenode) and 2,9 and 4,9 µg sheared gDNA, respectively, was used for library preparation. Size selection was done with the BluePippin device and a cut-off at 8,5 kb. The size-selected libraries were prepared for loading following the instructions generated by the SMRT Link software (PacBio, version 10) and the ‘HiFi Reads’ application. HiFi libraries were sequenced on 8M SMRT™ Cells for 30 hours with the Sequel^®^ II Binding 2.2 and the Sequel^®^ II Sequencing 2.0 chemistry on the Sequel^®^ II.

### PacBio library preparation and genome sequencing

Ultra-long gDNA of *Chelodina, Erymnochelys, Hydromedusa, Pelomedusa, Peltocephalus,* and *Podocnemis* was used for 10x genomic linked read sequencing following the manufacturer’s instructions (10X genomics Chromium Reagent Kit v2, revision B). To this end, 1 ng of ultralong gDNA was loaded into 10X genome GEM droplets (Gel Bead-In-EMulsions) using the Chromium device. gDNA molecules were amplified in individual GEMs using primers containing a specific 16 bp 10x barcode and the Illumina R1 sequence. After breaking the emulsions, pooled amplified barcoded fragments were purified, enriched, used for Illumina sequencing library preparation and sequenced on a NovaSeq 6000 S4 flow cell (2x150 paired-end).

### Bionano optical mapping

For *Chelodina, Erymnochelys, Hydromedusa, Pelomedusa, Peltocephalus* and *Podocnemis,* optical mapping was done with the Bionano Prep Direct Label and Stain DLS DNA Kit (catalog #8005, Bionano Genomics, San Diego), according to the manufacturer’s Protocol (Document Number 30206, Document Revision F). Briefly, 750 ng of ultra-long gDNA was fluorescently labeled at defined sequences, making use of the nicking-free Bionano Direct Label Enzyme (DLE-1). For further visualization, the DLE-1 labeled gDNA backbone was stained with DL-Green. Labeled molecules were imaged using the Bionano Saphyr system. Data were generated from one Bionano flow cell each.

### Chromatin conformation capturing (Hi-C)

Chromatin conformation capture (Hi-C) was performed using the ARIMA-HiC High Coverage kit according to the Arima documents (User Guide for User Guide Nucleated Blood, part number A160127 v00). Briefly, 50 µl of flash-frozen nucleated blood preserved in Ethanol of *C. parkeri, E. novaeguineae, E. madagascariensis, H. tectifera, P. somalica, P. dumerilianus,* and *P. unifilis,* were chemically crosslinked. The crosslinked genomic DNA was digested with a cocktail of four restriction enzymes and 5’-overhangs were filled in and labelled with biotin. Spatially proximal digested DNA ends were ligated, and the ligated biotin-containing fragments were enriched and used for Illumina library preparation according to the ARIMA user guide for library preparation using the Kapa Hyper Prep kit (ARIMA document part number A160139 v00). The barcoded Hi-C libraries were run on S2 and S4 flow cells of an Illumina NovaSeq 6000 at 2x100 or 2x150 cycles.

### Genome assembly

To assemble genomes from PacBio CLR, 10x linked-reads, Bionano optical maps and Arima Hi-C long-range interaction data, we followed the Vertebrate Genomes Project v1.6 pipeline ^43^ with modifications for error correction. We assembled the initial set of contigs using Falcon v1.8.1 and Falcon-Unzip v1.3.7 ^168^ and then removed any retained haplotigs from the primary assembly using purge-dups v1.2.3 ^169^. We then mapped 10x linked-reads to the assembled contigs using Longranger v2.2.2 and performed scaffolding with Scaff10x v4.2 (https://github.com/wtsi-hpag/Scaff10X). The 10x reads were then mapped to the scaffolds again using Longranger and spurious scaffolds were identified and broken using Break10x v4.2 (https://github.com/wtsi-hpag/Scaff10X). We then further scaffolded using the optical maps from Bionano and the Bionano Solve v1.6.1 “hybrid scaffold” tool (https://bionano.com/software-downloads/) and subsequently with Hi-C interaction data using Salsa2 v2.3 ^170^. For Hi-C read data mapping and processing we followed the VGP pipeline for mapping, filtering, deduplication and conversion to bed format for input to salsa2 (https://github.com/VGP/vgp-assembly/blob/master/pipeline/salsa/arima_mapping_pipeline.sh). Finally, we performed manual inspection of the Hi-C data mapped to assembled scaffolds using HiGlass v1.11.7 ^171^, correcting any mis-joins or missed-joins in the assembly by reordering and reorienting the sequences in the assembly.

Following scaffolding, we performed gap closing using the PacBio CLR long reads by mapping the reads to the genome using pbmm2 v1.4.0 and closed gaps using the polishing tool gcpp v2.0.2 (https://git.mpi-cbg.de/assembly/programs/gap_closing). We then corrected any base errors in the assembly by mapping the 10x Genomics linked-reads to the curated assembly using Longranger v2.2.2. We called variants using freebayes v1.3.2 with the argument “-g 600”. We then filtered the variants using bcftools view v1.12) with argument: ’-i ‘QUAL>1 && (GT=“AA” || GT=“Aa”)’ and consensus called using bcftools consensus with argument: ‘-Hla’. Finally we phased the chromosomes by applying an adapted version of the DipAsm pipeline https://doi.org/10.1038/s41587-020-0711-0, calling heterozygous sites using 10x reads and creating phase blocks by linking heterozygous sites in 10x, Hi-C and PacBio reads using Hapcut2 (git commit 1ee1d58) and whatshap (v1.6). We then used the whatshap-tagged PacBio and 10x reads, binned into H0 and H1, and H0 and H2, to haplotype-polish the chromosomes. We ran gcpp using the combined H0 and H1 haplotagged PacBio CLR reads, followed by freebayes polishing using the combined H0 and H1 haplotagged 10x reads as above, with the corresponding H2 and H0 tagged reads for the second haplotype.

For *Elseya novaeguineae*, we produced a haplotype-specific assembly from HiFi reads using hifiasm v0.16.0, integrating HiC reads for phasing ^172^. Both haplotypes were subject to filtering procedure using Foreign Contamination Screen Tool (FCS and FCS-GX) ^173^. Contamination free assemblies were then polished to remove unambiguous heterozygous sites. Specifically, we mapped HiFi reads back to each assembly with minimap2 v.2.21 ^174^ and called variants using DeepVariant v1.2.0 ^175^. We then filtered for sites with genotype 1/1 and a ‘PASS’ filter value, meaning that all or nearly all reads support an alternative sequence at this position and passed DeepVariants internal filters. Finally, we corrected corresponding nucleotide sites in the assembly using bcftools consensus v.1.12 ^176^.

To scaffold the polished haplotype assemblies of vampire bats, we first mapped the Arima HiC reads to each of the two assemblies independently using bwa-mem2 v.2.2 ^177^ and then used YaHS v1.1a ^178^. Finally, we performed manual curation of the two scaffolded haplotypes jointly analyzing them in PretextView v.0.2.5 (https://github.com/wtsi-hpag/PretextView). To this end, we remapped the HiC reads to both haplotype assemblies concatenated together using again bwa-mem2 but allowing for multi-mapping reads (-q 0) to avoid discarding information in regions identical between two haplotypes. We also identified telomeric sequences with tidk v.0.2.31 (https://github.com/tolkit/telomeric-identifier) and where necessary corrected wrong contigs orientations to have telomeres in the ends of resulting scaffolds. To finalize changes made via manual dual haplotype curation, as recently proposed by the Vertebrate Genome and Darwin Tree of Life Project, we used the rapid curation framework (https://gitlab.com/wtsi-grit/rapid-curation/-/tree/main) from the Genome Reference Informatics Team ^179^.

### Repeat masking

We modelled and masked repeats in 30 available turtle genome assemblies (Supplementary Table S5) ^43,45,58,59,69,78,180–183^ using RepeatModeler v2.0.1 and RepeatMasker v.4.1.2, respectively ^184,185^. Additionally, low complexity repeats were masked with WindowMasker v1.0 ^186^ and tandem repeats finder (trf) v4.09 was used to identify and mask tandem repeats with a maximum length of 3 Mbp ^187^.

### Genome alignments

We aligned each of the available turtle genome assemblies as well as our newly sequenced turtle genomes to the Red-Eared Slider Turtle *Trachemys scripta elegans* genome using a previously-established pipeline ^188^. We used LASTZ v.1.04.15 ^189^ with parameters (K = 2400, L = 3000, Y = 9400, H = 2000, and the LASTZ default scoring matrix) that have a high alignment sensitivity ^190^ to obtain local alignments that were then chained using axtChain ^191^ with default parameters except for linearGap=loose. We used RepeatFiller ^192^ with default parameters to add missed repeat-overlapping local alignments to the alignment chains and chainCleaner ^193^(default parameters except for minBrokenChainScore = 75,000 and -doPairs) to improve alignment specificity.

Additionally, we aligned our newly sequenced genomes to genome assemblies of four additional turtles (*Chelodina mydas*, *Gopherus evgoodei*, *Mauremys reevesii*, *Dermochelys coriacea*) as well as chicken (*Gallus gallus domesticus*) and Australian Saltwater Crocodile (*Crocodylus porosus*). The resulting alignments were used for genome annotation and orthology inference with TOGA. For stem-turtle gene-loss analyses, we also aligned 7 Cryptodira genomes (*Chelonia mydas*, *Gopherus evgoodei*, *Mauremys reevesii*, *Dermochelys coriacea*, *Rafetus swinhoei, Carettochelys insculpta,* and *Trachemys scripta elegans*) to genome assemblies of the chicken and the Australian Saltwater Crocodile. Genome alignments of turtles to the more distantly related chicken and crocodile, were run with even more sensitive LASTZ parameters (setting L=2800, K=2200). All assembly accessions are listed in Table S5.

### Genome annotation and orthology inference

We used TOGA v.1.0 (Tool to infer Orthologs from Genome Alignments) to annotate protein coding genes and infer orthologues for phylogenomic- and gene-loss analyses (https://github.com/hillerlab/TOGA). TOGA uses pairwise whole genome alignments of an un-annotated query genome to an annotated reference genome and a machine learning classifier to annotate protein-coding genes and infer orthology, i.e., distinguish orthologous from paralogous or processed pseudogene loci. ^47^.

To maximize genome annotation for the newly sequenced turtle genomes, we used multiple reference species we used and integrated results from seven references: Green Sea Turtle *Chelodina mydas* (GCF_015237465.1), Goodes Thornscrub Tortoise *Gopherus evgoodei* (GCF_007399415.2), Red-Eared Slider Turtle *Trachemys scripta elegans* (GCF_013100865.1), Reeves’s Turtle *Mauremys reevesii* (GCF_016161935.1), Leatherback Sea Turtle *Dermochelys coriacea* (GCF_009764565.2), chicken *Gallus gallus domesticus* (GCF_000002315.5) and Australian Saltwater Crocodile *Crocodylus porosus* (GCF_001723895.1).

To assess the completeness of the new turtle genomes and their annotations, we ran BUSCO v. 5.2.2 (--mode ‘protein’) on the combined TOGA annotations using the sauropsida_odb10 dataset with 7,840 BUSCO genes ^48^. Comprehensive annotations often include multiple isoforms per gene, so duplicated BUSCO genes do not indicate a problem, but rather a comprehensive transcript annotation. Accordingly, we focus on the number of complete and incomplete (missing and fragmented) genes (Figure 1B).

### Exon-by-exon gene alignments

For phylogenomic analyses, we used the longest isoforms of one-to-one orthologs of the query species and *T. scripta elegans* that were present in at least 80% of the query species. In cases of multiple longest (equal length) transcripts for the same gene, we selected those present and intact in the greatest number of assemblies. Codons with frameshifting insertions or deletions and premature stop codons were masked with ‘NNN’ to maintain the reading frame. Subsequently, we generated codon-aware exon-by-exon multiple sequence alignments using MACSE2.0 ^194^ and removed poorly aligning regions with HmmCleaner v.0.180750 using the default parameters ^195^.

### Phylogenomics

To reconstruct the turtle phylogeny from genome-wide data, we used whole-gene codon alignments and estimated a species tree using ASTRAL v.5.7.1 ^196,197^. We used RAxML v.8.1.16 to compute gene trees using three independent replicates, a GTR + GAMMA model and a rapid-hill-climbing algorithm ^198^. The resulting gene trees were used as input for ASTRAL to compute a species tree. We used 100 bootstrap replicates to assess node supports of the species tree and calibrated it using the congruify.phylo function implemented in geiger package ^199^ and the most complete turtle timetree ^42^.

### Positive selection

To identify genes evolving under episodic positive selection in stem turtles, we used our whole-gene codon alignments using our inferred phylogenetic tree topology as input and ran the adaptive branch-site random-effects likelihood method aBSREL ^200^ implemented in HYPHY v.2.5.8. aBSREL uses the ratio of non-synonymous to synonymous substitution rates (dN/dS) to infer whether sites evolved under purifying selection (dN/dS < 1), neutrally (dN/dS ≈ 1) or under positive diversifying selection (dN/dS > 1). We ran aBSREL in the foreground mode on the branch leading to all extant turtles and enabled synonymous rate variation to improve the rate of false positives. In total, we screened 11,141 one to one orthologous genes for selection. Alignments of all genes under significant positive selection (p<0.01) that are mentioned in this manuscript were further manually inspected to avoid misalignments generating spurious results.

### Population history inference

We mapped 10X reads to published genomes of *Mesoclemmys tuberculata* (GCA_007922155.1), *Podocnemis expansa* (GCA_007922195.1), *Emydura subglobosa* (GCA_007922225.1) and *Pelusios castaneus* (GCA_007922175.1), as well as all newly generated Pleurodira reference genomes using Longranger (v2.2.2). Specifically, we grouped reads based on their 10X barcodes, trimmed barcodes, conducted error-correction, sorted and aligned the reads to the genomes. The resulting mappings were quality adjusted (mpileup -C50), and sites with less than one third or more than twice the average genome wide coverage were removed with bcftools (v1.10.2) ^202^. Mean coverages ranged from 48X to 85X (*E. subglobosa*: 48X, *M. tuberculata*: 54X, *P. castaneus*: 52X, *P. unifilis*: 85X, *P. expansa*: 48X, *H. tectifera*: 76X, *C. parkeri*: 128X, *E. novaeguineae*: 28X, *E. madagascariensis*: 44X, *P. dumerilianus*: 64X, *P. somalica*: 121X). Finally, we converted variant calls (including invariant sites) to fasta sequences and the PSMC-specific input format with fq2psmcfa (-q20) (https://github.com/lh3/psmc).

We initially ran PSMC with the settings “-N25 -t15 -r5”, varying the parameter patterns (-p) between “4+25*2+4+6”, “2+2+25*2+4+6” and “1+1+1+1+25*2+4+6”. To increase the resolution of population size inferences, while avoiding overfitting, we increased the number of atomic time intervals and chose the setting with the highest number of windows that achieved at least 10 recombination events for each time window after 20 iterations. PSMC inferences were scaled using a mutation rate of 7.9 × 10^−9^ ^203,204^ and generation times of 15 years for *Emydura subglobosa, Elseya novaeguineae* and *Pelusios castaneus*, 25 years for *Podocnemis unifilis*, *Hydromedusa tectifera*, *Mesoclemmys tuberculata*, *Peltocephalus dumerilianus, Pelomedusa somalica, Erymnochelys madagascariensis, Chelodina parkeri* and 50 years for *Podocnemis expansa*.

### Linking population histories to historical climate, biomes and sex determination

We used generalized linear mixed models (GLMM) to investigate the drivers of side-necked turtle population declines inferred with PSMC across the last 140k years. More specifically, we investigated the influence of historical climate change and ecosystem change and whether the effect differed for turtles with genetic and temperature dependent sex determination.

To assess whether historical biome shifts contributed to the inferred turtle population declines, we first georeferenced turtle distribution ranges ^63^ in QGIS ^205^. We extracted current and historical biomes across the last 140,000 years from Allen et al. (2020), who simulated climate-driven biome shifts with a Dynamic Global Vegetation Model. Climate forcing was provided by the HadCM3 climate model and included rapid climate fluctuations known as“Heinrich Events” ^62^. Georeferenced distribution areas were digitized using polygons and the data extracted using “Selection by Location” in QGIS. Using current biomes, a biome was classified as suitable for a species, if it made up at least 10% of the current distribution range. For each time point with historical biome data, biome suitability was calculated as the proportion of the current distribution range covered by suitable biomes.

Recent and past climatic data (monthly mean temperature, mean monthly precipitation total and monthly mean ‘cloudiness’) were those used by ^62^ to simulate the biomes. Recent climatic data were sourced from the CRU TS 3.0 dataset ^60^ at half-degree spatial resolution, with climatic values for shelf grid cells exposed by the lowered global sea level at the Last Glacial Maximum (LGM) estimated using thin-plate spline surfaces fitted to data for land cells in the CRU dataset ^206^. Palaeoclimatic data for each time slice were obtained by applying to the recent data anomalies computed between the palaeoclimate simulation for that time slice and a pre-industrial climate simulation. Anomalies were additive for temperature and multiplicative for precipitation and ‘cloudiness’. Palaeoclimate simulations were made using the HadCM3 fully-coupled atmosphere–ocean general circulation model with boundary conditions and forcings as used in the experiments by ^61^. However, unlike ^61^, our simulations used HadCM3 coupled to TRIFFID (Cox, 2001)[BH5], a simple dynamic global vegetation model, thus dynamically adjusting the land surface characteristics to the simulated climate. In addition, pre-LGM ice-sheet configurations were specified using a different and likely more realistic approach, as described by ^207^.

To integrate biome and climate data with psmc-inferred historical population sizes, we calculated the means for all data points within the corresponding psmc time window. Since PSMC inferences for the first time window are less reliable, we excluded the first window from each time series. We also tested alternative filtering of the PSMC input data, which did not affect the general results. More specifically, we also tried excluding all time windows with midpoints <20 kya or <10 kya and excluding the last PSMC time window, if no biome data was available for >50% of the time spanned by this window. We then fitted 615 generalized linear mixed models using the MCMCglmm R package (v. 2.34) ^208^. These models all had log-transformed population size as the response variable and differed in their fixed effects (explanatory variables), their random effects, and whether they were phylogenetically constrained. More precisely, the fixed effects component of the models ranged from only an intercept (an empty model that serves as a baseline) to all possible combinations of the following explanatory variables: a) biome suitability, b) type of sex determination, c) mean monthly ambient temperature, d) ambient temperature range (i.e., the difference between the maximum and minimum monthly temperatures), and e) time—to account for declining population sizes towards the present due to population structure ^64^. Some models also allowed for interactions between i) type of sex determination and mean ambient temperature, ii) type of sex determination and ambient temperature range, and iii) biome suitability and time. Regarding random effects, we again specified all possible model variants, i.e., with no random effects and with (phylogenetic or non-phylogenetic) random effects of species identity on the intercept and/or the slopes of the explanatory variables per model. The priors that we specified were relatively uninformative, namely the default Gaussian prior for fixed effects, parameter-expanded priors for the random effects (where applicable), and an inverse-gamma prior for the residual variance. For each candidate model, we executed four independent chains for 60 million generations, with samples from the posterior being recorded every 2,500 generations after the first 6 million generations, which we discarded as burn-in. To ensure that the four chains per model had converged on statistically equivalent posterior distributions and that each parameter was sufficiently sampled, we verified that each parameter had an effective sample size of at least 1,000 (in each chain) and a potential scale reduction factor less than 1.1. To identify the best-fitting model across our catalogue, we first removed models with one or more fixed effects (excluding the intercept and main effects of interactions) whose 95% highest posterior density interval included zero, and then compared the remaining models based on their Deviance Information Criterion values (DIC; averaged across the four independent chains) and the corresponding DIC weights. Finally, for the best-fitting model, we calculated the marginal and conditional coefficients of determination, *R*^2^_m_ and *R*^2^_c_ ^209^. These, respectively, represent the fraction of variance explained by the fixed effects or by both the fixed and the random effects.

### Y-chromosome microdissection and sequencing

Mitotic chromosome suspensions from a male *Elseya novaeguineae* were prepared from whole blood cell culture (for detailed protocol see Mazzoleni et al. 2020). Chromosome spreads were stained by C-banding ^210^ to identify the Y chromosome, which has a prominent C-positive heterochromatic band, missing in the X chromosome. We microdissected 20 Y chromosomes, in two replicates, by sterile glass needles using a Zeiss Axiovert S200 inverted microscope (Oberkochen, Germany) equipped with an Eppendorf TransferMan NK2 mechanical micromanipulator (Hamburg, Germany). The microdissected chromosomal material was amplified by a commercial kit (WGA4 kit, Sigma-Aldrich) and part of the amplified material was used for chromosome painting to verify that the microdissected material includes genomic regions from the desired chromosome (for a detailed protocol see ^211^). In situ hybridization images were captured by an Olympus BX51 fluorescence microscope equipped with an Olympus DP70 digital camera. The rest of the amplified microdissected material was sequenced in Illumina platform with 150 base pairs (bp) pair-end option (DNA-seq) (Novogene, UK). Finally, sequencing reads were mapped to our *Elseya novaeguineae* reference genome and with bwa-mem ^212^ and mean coverage per chromosome was estimated with samtools v.1.14 ^213^ to identify the sex chromosome.

### Sex chromosome identification and evolution

We used Findzx v.1 to identify sex chromosomes in Chelidae ^67^. To this end, we used illumina technology to re-sequence one male and one female of *Chelodina parkeri*, two males and three females of *Hydromedusa tectifera*, and one male and one female of *Elseya novaeguineae*. Sequencing adapters were trimmed with trimmomatic (Parameters: LEADING:3, TRAILING:3, SLIDINGWINDOW:4:15, MINLEN:36”) and reads subsampled to the same coverage (30.6 X) for all samples ^214^ mapping with bwa-mem ^212^, and variants were called and analyzed as implemented in Findzx v.1 ^67^. Using the same approach, we confirmed results for potential micro-cex chromosomes in *H. tectifera* and *C. parkeri* by mapping reads from these to species against the *E. novaeguineae* reference genome. To distinguish between X and Y in the haplotype-resolved assembly of *Elseya novaeguineae*, we extracted and estimated the length of GATA-motifs that were masked by repeat masker from chromosome-size scaffolds of both haplotypes. To further explore protein coding differences between the sex chromosomes, we collected genes with an intact projection on the *Elseya novaeguineae* sex chromosome that were not intact in the other haplotype.

Since fusions between sex chromosomes and autosomes can resolve intralocus sexual antagonism we estimated the likelihood of the observed sex chromosome autosome fusion in Chelidae given the karyotype, the sex determination system and the total number of fusions based on ^79^.

### Chromosome evolution reconstruction

To gain insight into turtle chromosome evolution, we reconstructed ancestral chromosomes using AGORA ^68^. To reconstruct chromosome-size contiguous ancestral regions, we used our newly assembled chromosome-level Pleurodira genomes and chose the 7 best Cryptodira genome assemblies that also covered karyotype diversity within Cryptodira ^26^ as well as chicken and crocodile.

We subsequently trimmed our newly created genome-wide turtle phylogeny to include only the selected species. Orthology groups were created for each ancestral node in the phylogeny, by selecting genes that were one-to-one orthologs and never lost or missing across any of the descending species based on TOGA classification (I, PI, UL) using *T. scripta elegans* as reference. Similarly, gene order files for each species were generated for each species using one-to-one orthologs projected from *T. scripta elegans*. With these input files, AGORA was run in the generic multi pass mode to allow automatic selection of the best settings (agora-generic.py). For curation and visualization, resulting ancestral and extant genomes were plotted and compared using syntenyPlotteR ^215^.

Subsequently, we manually curated CARs (Figure S13). We corrected over splitting of ancestral chromosomes due to rearrangements for which AGORA cannot resolve the ancestral orientation, resulting in splitting of CARs. If such CARs were adjacent in both descendent lineages, we re-named them and treated them as a single ancestral chromosome in subsequent analyses (CAR_X, CAR_Y → CAR_XA CAR_XB). Similarly, CARs were fused, if they were adjacent in one descending lineage and in the most closely related outgroup. To focus on chromosome-size CARs, we removed CARS containing less than 20 genes, unless they were the largest representative of a chromosome in a descending lineage. Finally, to account for potentially missing chromosome size representation of chromosomes in assemblies of extant species, we assumed a CAR was falsely missing in an ancestor, if it was present in one descending lineage and the most closely related outgroup. We started the curation at genome assemblies of extant species followed by increasingly older ancestral nodes so that already curated genomes would serve as references for further curation of their ancestors. For extant species, chromosome numbers were mostly known and corresponded well to the published, chromosome level scaffolds. However, we identified discrepancies for the softshell turtle *Rafetus swinhoei* for which only 25 scaffolds have been assigned chromosome level following our manual curation, we ended up with a most likely karyotype of 2n=64-66 chromosomes. Since 2n=64-66 is a conserved karyotype among other softshell turtles and 2n=66 is likely the most common ^26,216^, we base our analyses on this karyotype. Additionally, the genome assembly of *Gopherus evgoodei* initially only contained 1n=24 chromosome-level scaffolds. We found that the chromosome homologous to NC_052644 in *Mauremys reeversi* was fragmented (NW_022060016 and NW_022060027). Similarly, the chromosome homologous to NC_052636 in *Mauremys reeversi* was fragmented (NW_022060064, NW_022060072, NW_022060107, NW_022060109). Taking this into account, we ended up with 2n=52 chromosomes for *Gopherus evgoodei*, which is further in agreement with analyses indicating 2n=52 chromosomes in *Gopherus* ^69^. Chromosome fission and fusion rates were calculated by dividing the number of the respective events by the branch length. To assess whether chromosome mutation rates were increased in focal lineages, we calculated poisson probabilities for observing at least the reconstructed number of events given the branch lengths and background chromosome mutation rate across the entire dataset. P-values were adjusted for multiple testing with Bonferroni correction.

### Chromosome breakpoint repeat content

To investigate whether repeat content promotes chromosome fissions in turtles, we focused on fissions in the only branch with a significantly increased chromosome fission rate. We excluded one recurrent breakpoint that also occurred in other lineages. For each remaining fission, we determined the neighboring genes on either side of the breakpoint in the most recent common ancestor before the fission occurred. We then used these genes to determine breakpoint regions in extant genomes. To keep inferences comparable across lineages, we filtered genome annotations of extant species to include only genes present in the most recent reconstructed ancestor before the fission occurred. Using these filtered annotations, we extracted breakpoint region features for subsequent analyses in extant query genomes by i) selecting the region between the upstream breakpoint gene and its next downstream neighboring gene, ii) selecting the region between the downstream breakpoint gene and its next upstream neighboring gene and iii) calculating the average of i) and ii) (Figure S14). If the upstream/downstream breakpoint gene was the last/first on a chromosome, we considered the region between that gene and the end/start of the scaffold. To compare breakpoint region features to the genome wide background, we sampled 100 random neighboring gene pairs in the most recent common ancestor before the chromosome fission occurred. To calculate the length/proportion of repeats, we combined simple repeats and repetitive sequences masked by repeat masker using bedtools merge v2.28.0 ^217^. If a region did not contain any repeats, it was excluded from the analyses for the respective species. For clades with multiple extant species, we summarized statistics by calculating mean repeat contents and mean lengths for each breakpoint- and background region. A Shapiro-Wilk-Test was used to test for normality and a paired T-Test was used to evaluate whether repeat content and length of breakpoint regions differed significantly between clades.

### Gene losses in stem-turtles

To detect gene losses in stem turtles, we used the TOGA classifications of gene intactness across the reference genomes of 14 turtle species (7 Cryptodira and 7 Pleurodira) using both crocodile and chicken as references. We considered a genes to be likely lost in the stem-turtle branch, if it met the following five criteria: It was i) classified as lost, missing or uncertain loss in all 14 turtles using chicken as reference, ii) classified as lost, missing or uncertain loss in all 14 turtles using crocodile as reference, iii) classified as lost in at least one turtle using chicken as reference, iv) classified as lost in at least one turtle using chicken as reference and v) classified as either intact or partially intact in crocodile using chicken as the reference. Subsequently, all alignments of genomic regions and inactivating mutations were manually inspected to confirm losses of candidate genes.

To evaluate whether genes associated with disproportionate dwarfism were enriched among lost genes we used a hypergeometric test based on a total of 72 genes annotated with with the term “disproportionate dwarf” in the “Mammalian Phenotype Ontology Annotations” database (MGI 6.24; database update 07/22/2025) and 23,312 protein coding genes in the human genome (GCF_000001405.40-RS_2024_08) ^97,98^.

To explore whether genes lost in turtles were otherwise conserved across tetrapods, we investigated how often these genes were classified as lost (“L”) based on TOGA across placental mammals using the human as the reference species and across birds using the chicken as the reference species. To avoid artifacts due to limited assembly quality, we selected genomes with less than 1,900 gene losses resulting in a final dataset of 754 genomes of 651 placental mammals and 107 genomes of 107 birds.

## Acknowledgment

This work was supported by the LOEWE-Centre for Translational Biodiversity Genomics (TBG) funded by the Hessen State Ministry of Higher Education, Research and the Arts (LOEWE/1/10/519/03/03.001(0014)/52), the Johanna Quandt Young Academy (JQYA Fellowship to L.H.), grants from German Research Foundation (L.H.: HI2214/1-1, Project number: 530763738, M.H.: HI1423/6-1), Michail Rovatsos was supported by the Czech Science Foundation (project GAČR 23-07347S). We are grateful to Carola Greve, Alexander Ben Hamadou and Charlotte Gerheim for assistance in the lab and to Bradley Shaffer lab for generating genomes of *M. tuberculata*, *P. expansa*, *P. castaneus*, and *E. subglobosa* and making the data freely available for analysis. Chromosome microdissection took place at the lab of Prof. Antonio Sanchez Baca at University of Jaen, Spain.

## Data and Code Availability

Genomes and raw data for *Pelomedusa somalica* (GCA_051311615.1), Hydromedusa tectifera (GCA_049999965.1) and Podocnemis unifilis (GCA_050000005.1) are available for download on NCBI and GenomeArk (https://www.genomeark.org/). The remaining genomes have been submitted to NCBI (accessions pending). TOGA annotations and ancestral genome reconstructuns are available at https://genome.senckenberg.de/download/TOGA/.

