## Supplement for "Side-necked turtle genomes reveal chromosomal dynamics, skeletal innovation and cancer resistance"

The supplement contains

- Figures S1 to S14

Supplementary tables S1-S5 are provided as sheets in an Excel file.

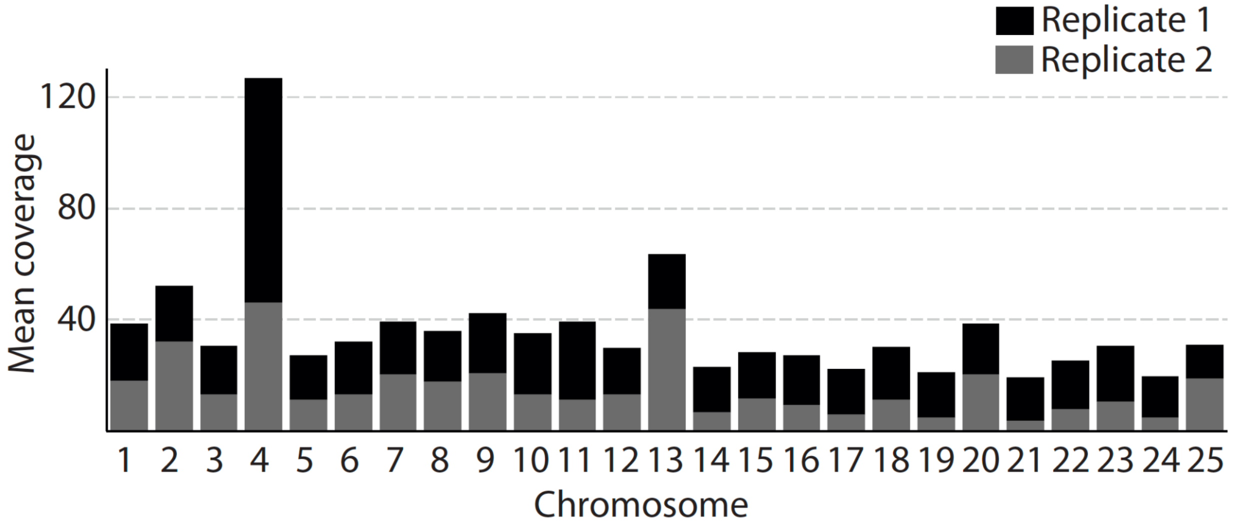


**Figure S1. Y chromosome capture for *Elseya novaeguineae*.** For each chromosome of the first haplotype of *Elseya novaeguineae*, the mean coverage of mapped chromosome capture sequencing data is shown. Two independent experiments (replicates) are plotted in black and grey, respectively. Similar to our bioinformatic analyses, chromosome capture further supports that chromosome four in our assembly represents the pair of sex chromosomes.

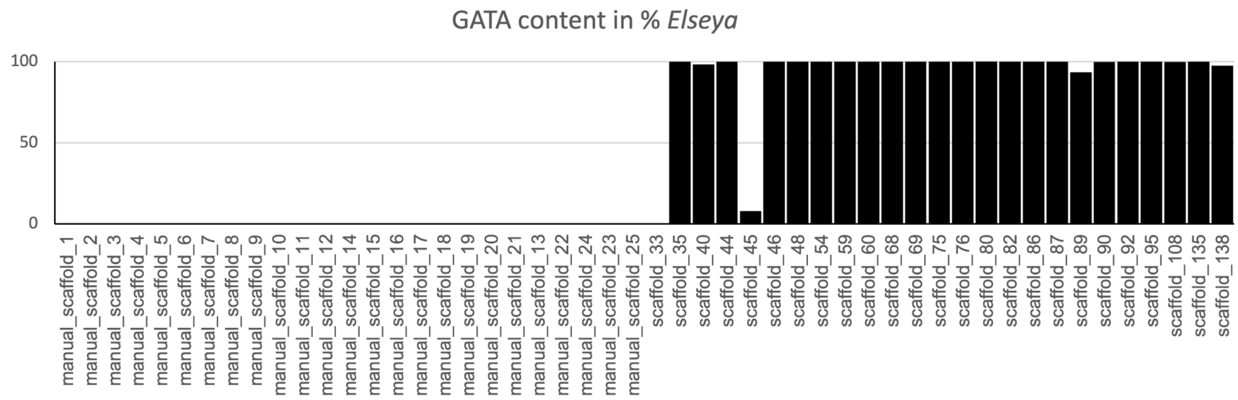


**Figure S2. GATA motif repeat content in haplotype 1 of *Elseya novaeguineae*.** For each scaffold, bars show the proportion (in %) made up by GATA repeat motifs. Several subchromosomal scaffolds consist entirely of GATA repeat motifs.

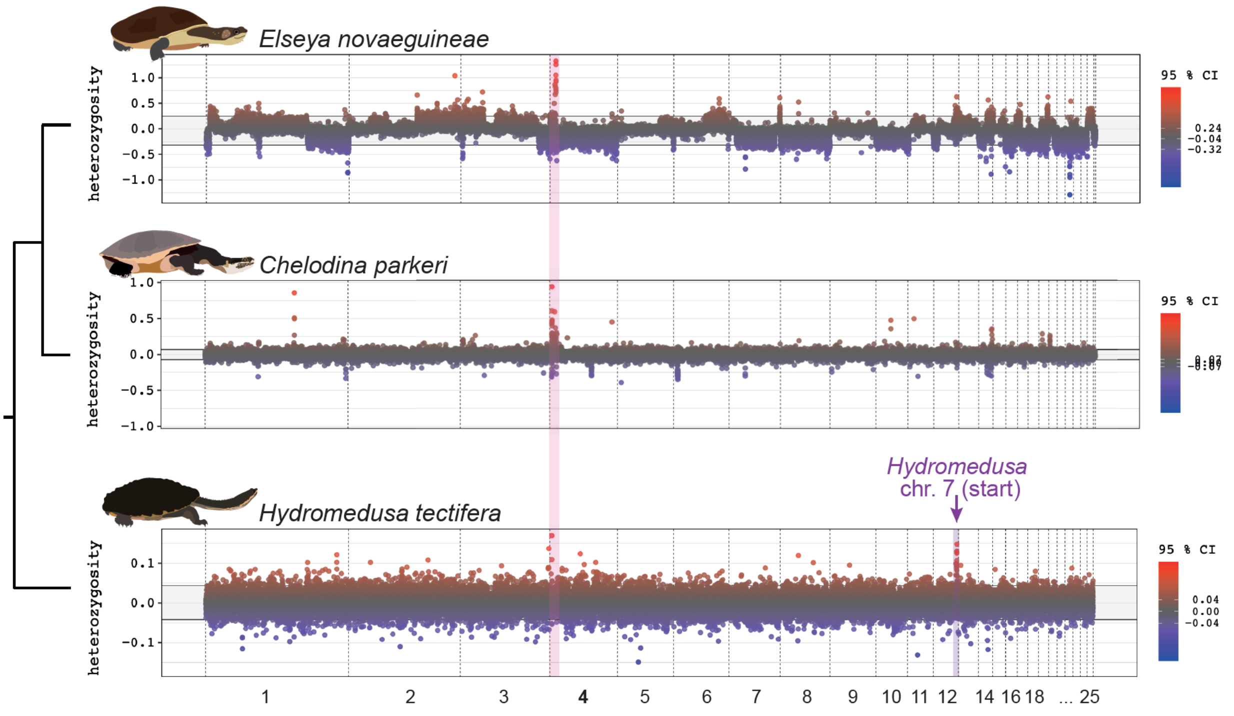


**Figure S3. Sex chromosome identification by mapping reads to the *Elseya* genome.** For each species, we ran the same pipeline as described in the main text, but mapped reads to *Elseya novaeguineae* (Hap 1) instead of reference genomes of each species. In all cases, the highest outlier window is located at the start of chromosome 4. Thus, our results provide strong support for an ancestral micro-sex and indicate that one homologous region contains sex linked-regions across all three species. Additionally, for *Hydromedusa tectifera* a different outlier region at the end of chromosome 12, which corresponds to the start of chromosome 7 in *Hydromedusa* might either reflect a more recent sex chromosome turnover in this species.

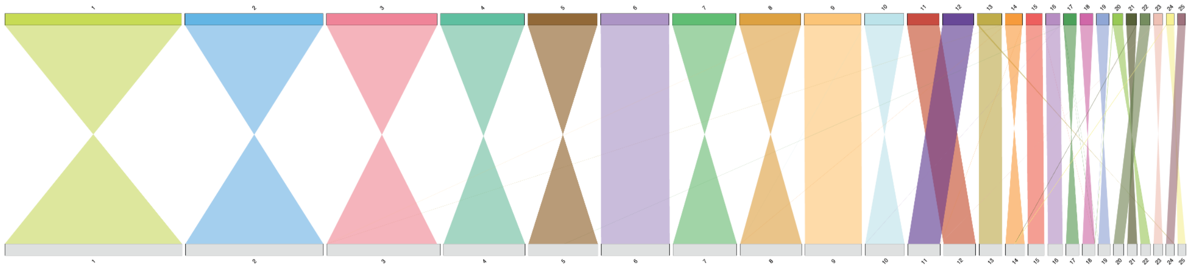

**Figure S4. Synteny between the two haplotypes of *Elseya novaeguineae*.** Fine-scale synteny between chromosome-size scaffolds of haplotype 1 (top) and haplotype 2 (bottom) is shown based on a whole genome alignment using 5kb alignment windows for illustration. Similar to the autosomes, chromosome 4 of each haplotype representing the neo-Y (top) and neo-X (bottom) chromosomes align from start to end. This further supports that that the ancestral micro-X chromosome was not lost when the micro-Y chromosome fused to a large autosome, but also fused to the same autosome in the ancestors of *Elseya*. Note that fully inverted chromosomes just represent arbitrary chromosome orientation in the assembly.

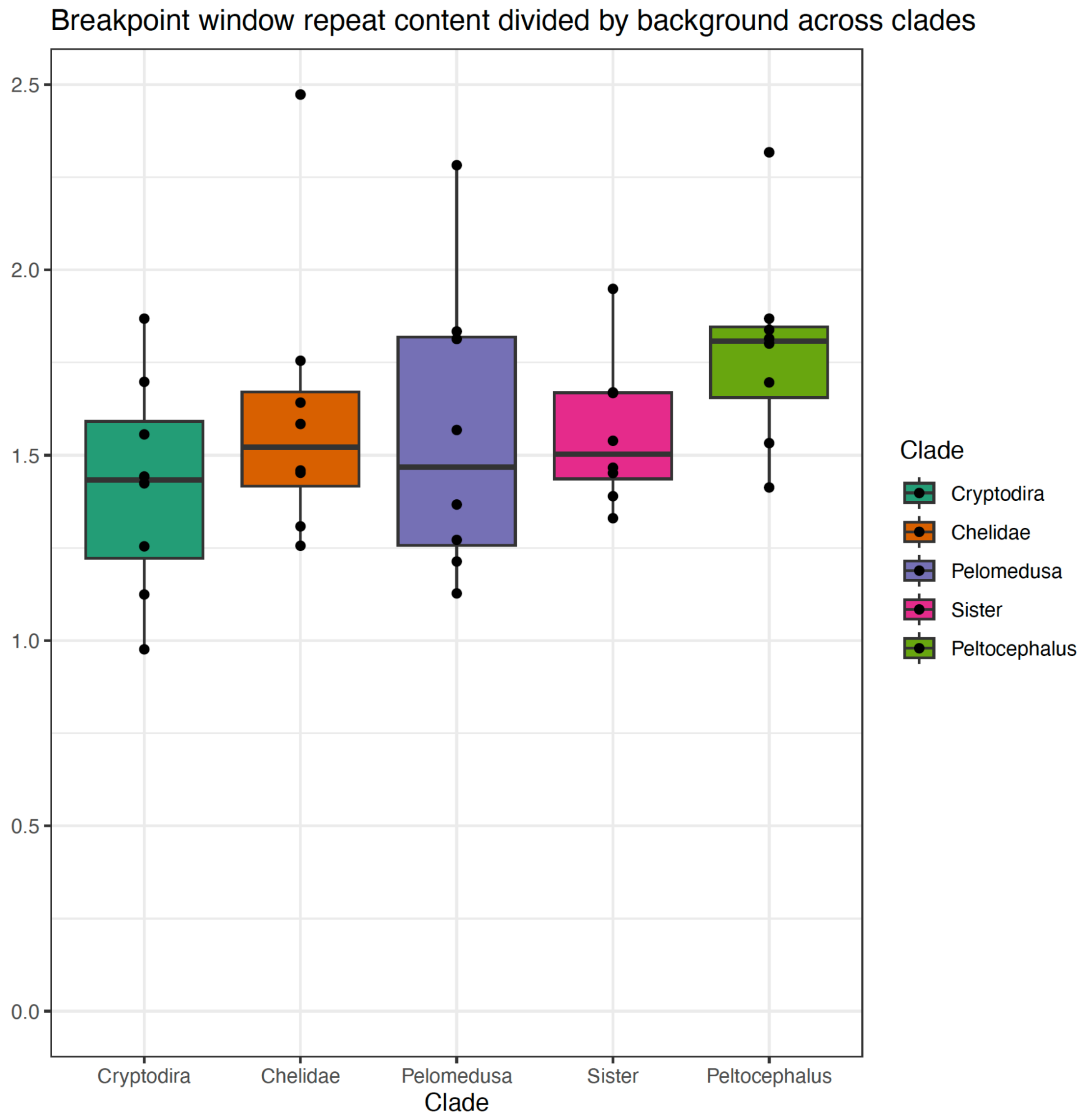


**Supplementary Figure S5. Proportion of breakpoint repeat content compared to genomic background.** For each clade the repeat content at the breakpoint window divided by the genomic background repeat content is shown. Aside from *Peltocephalus*, repeat contents at breakpoints remain roughly proportional to the genomic background.

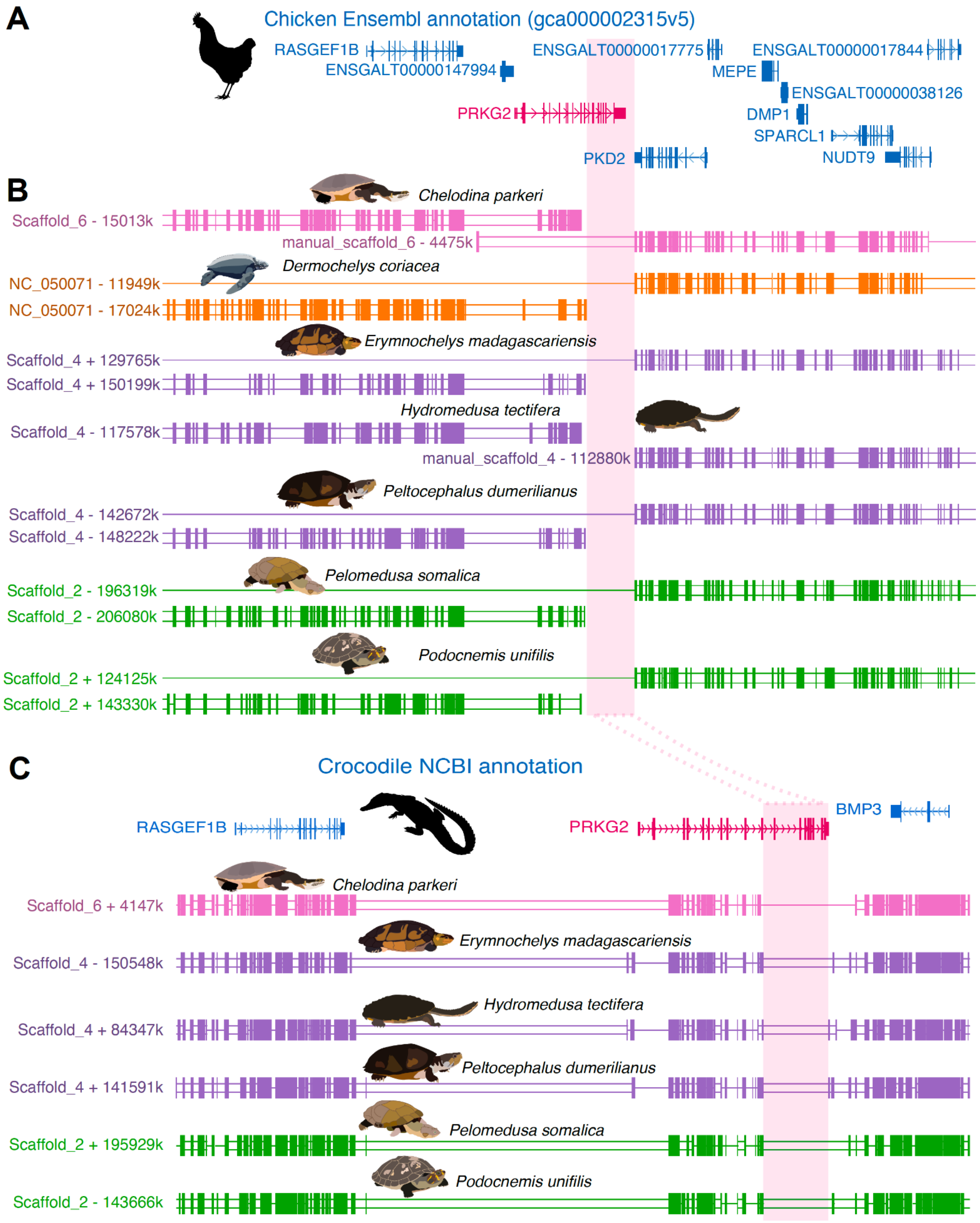


**Figure S6.** ***PRKG2* deletion in stem-turtles.** A) Gene order around the focal *PRKG2* (red) in the chicken. For clarity, only one transcript is shown per gene.

B) Alignment chains for seven high quality turtle genomes against the chicken show that *PRKG2*-loss coincides with deletion of the last seven exons in stem-turtles, which also overlaps an evolutionary breakpoint (highlighted in light red) of an intrachromosomal rearrangement in chicken.

C) Gene order around the focal *PRKG2* (red) in the saltwater crocodile (top). For clarity, only one transcript is shown per gene. Alignment chains of six turtles confirm both the deletion of seven exons and a shared *PRKG2*-loss.

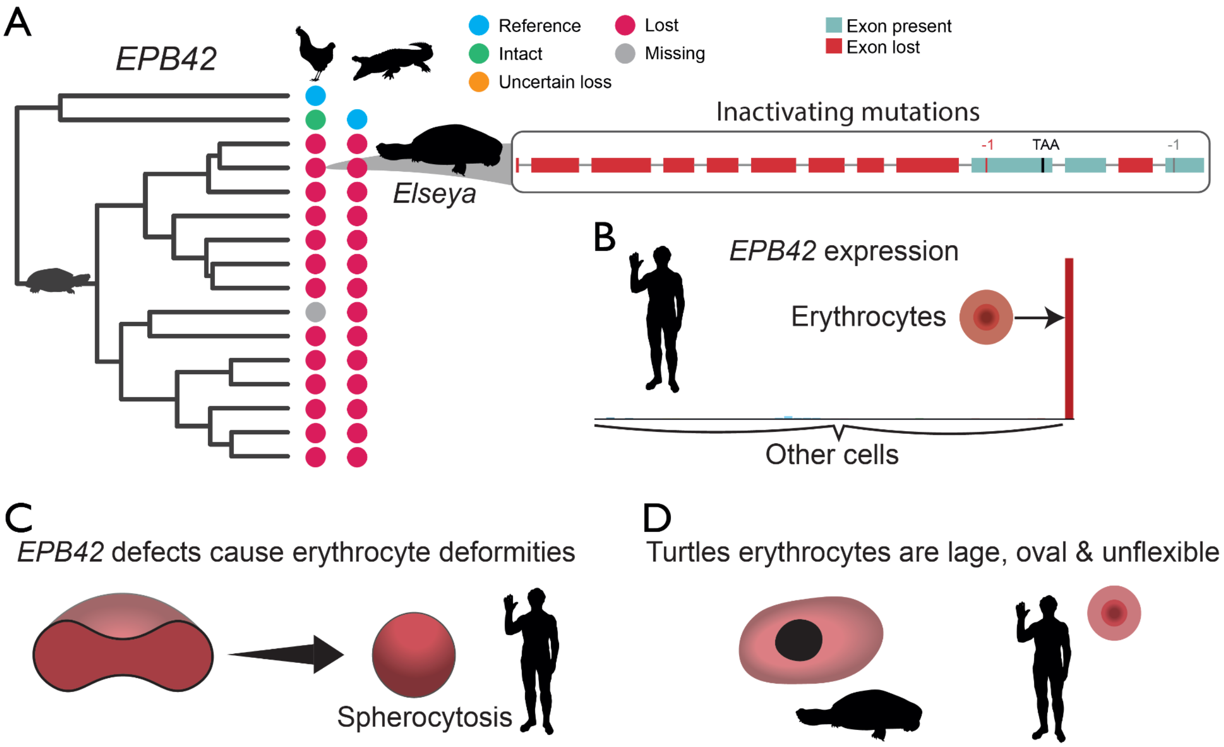


**Figure S7. *EPB42*-loss in turtles.** A) *EPB42* is lost in turtles. Using chicken and crocodile as references, colored circles tips show the gene loss pattern in 14 turtle genomes across the turtle phylogeny using chicken (left column) and crocodile (right column) as references. Inactivating mutations are shown in both isoforms for one representative side-necked and hidden neck turtle. B) *EPB42* has erythrocyte-specific gene expression in humans (https://www.proteinatlas.org/). C) In humans, *EPB42*-loss causes Spherocytosis (round Erythrocytes). D) Illustration of a turtle erythrocyte next to a human erythrocyte. Turtle erythrocytes are nucleated, very large and oval.

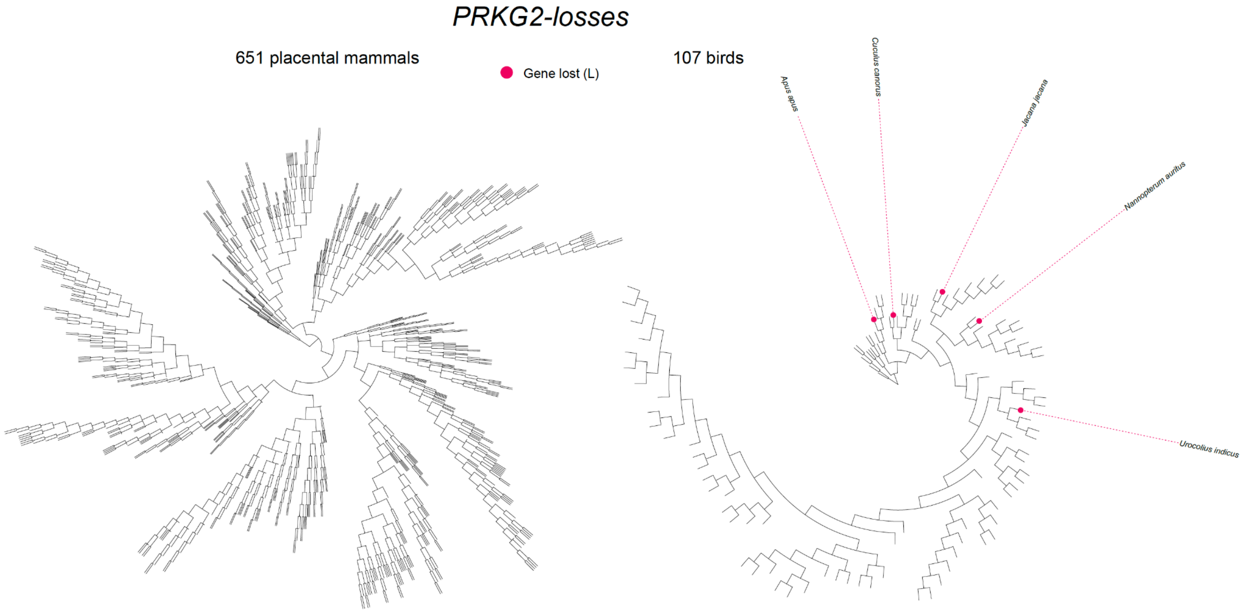


**Figure S8. *PRKG2*-losses in birds and placental mammals.** Red circles show gene losses (L) of *PRKG2* across a phylogeny including 754 high-quality genomes of 651 placental mammals (left) and across a phylogeny including 107 high-quality genomes of 107 birds (right). Species names are provided for genomes with a gene loss. High-quality genomes were chosen based on the abundance of inactivation mutations. Genomes with inactivating mutations in >1,900 genes were excluded from the analysis.

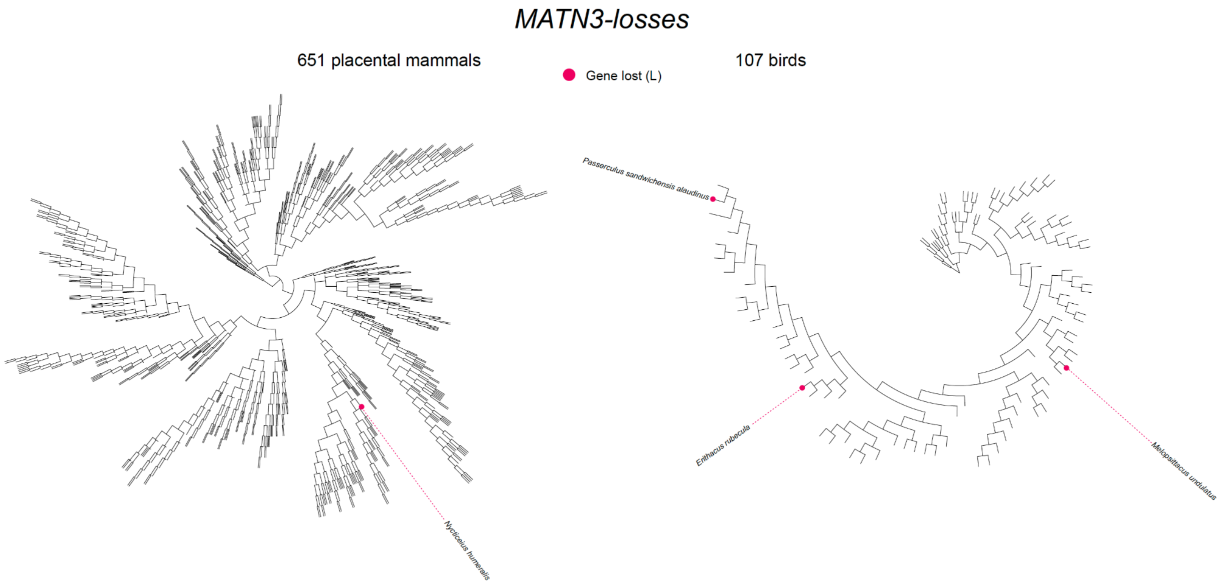


**Figure S9. *MATN3*-losses in birds and placental mammals.** Red circles show gene losses (L) of *MATN3* across a phylogeny including 754 high-quality genomes of 651 placental mammals (left) and across a phylogeny including 107 high-quality genomes of 107 birds (right). Species names are provided for genomes with a gene loss. High-quality genomes were chosen based on the abundance of inactivation mutations. Genomes with inactivating mutations in >1,900 genes were excluded from the analysis.

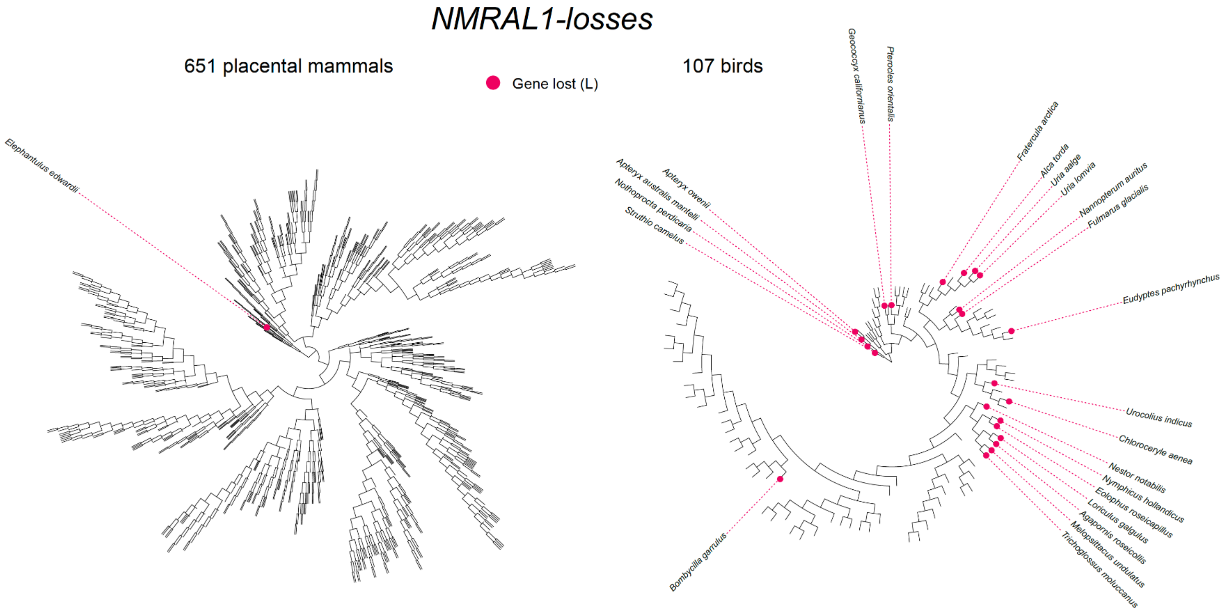


**Figure S10. *NMRAL1*-losses in birds and placental mammals.** Red circles show gene losses (L) of *NMRAL1* across a phylogeny including 754 high-quality genomes of 651 placental mammals (left) and across a phylogeny including 107 high-quality genomes of 107 birds (right). Species names are provided for genomes with a gene loss. High-quality genomes were chosen based on the abundance of inactivation mutations. Genomes with inactivating mutations in >1,900 genes were excluded from the analysis.

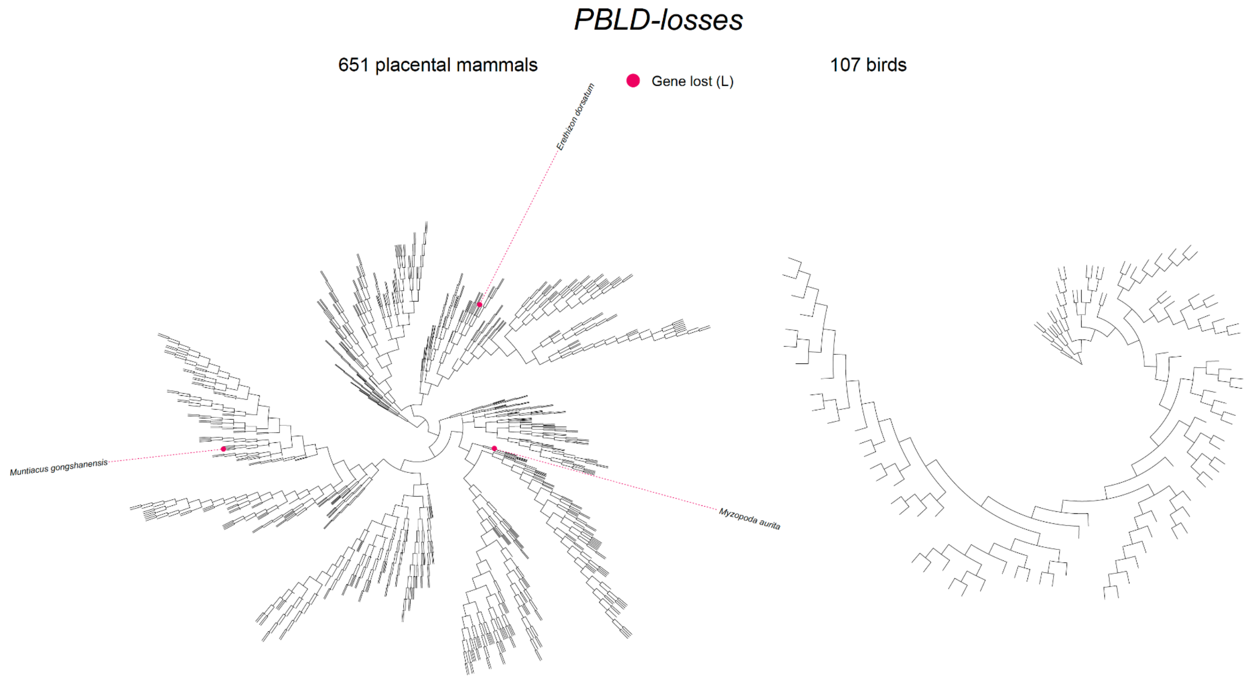


**Figure S11. *PBLD*-losses in birds and placental mammals.** Red circles show gene losses (L) of *PBLD* across a phylogeny including 754 high-quality genomes of 651 placental mammals (left) and across a phylogeny including 107 high-quality genomes of 107 birds (right). Species names are provided for genomes with a gene loss. High-quality genomes were chosen based on the abundance of inactivation mutations. Genomes with inactivating mutations in >1,900 genes were excluded from the analysis.

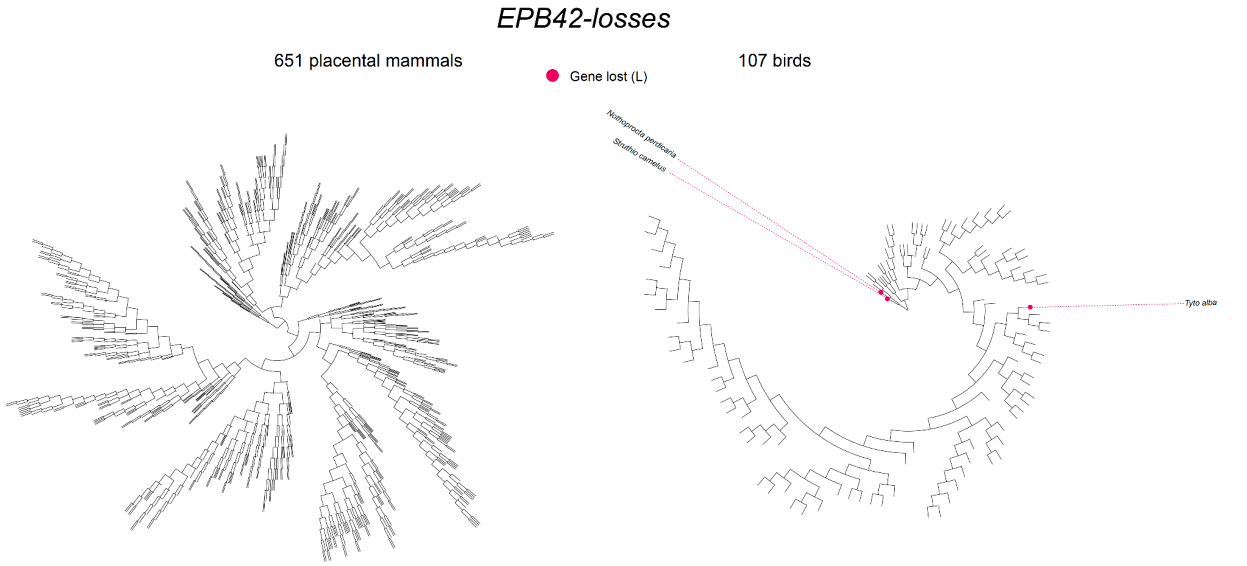


**Figure S12. *EPB42*-losses in birds and placental mammals.** Red circles show gene losses (L) of *EBP42* across a phylogeny including 754 high-quality genomes of 651 placental mammals (left) and across a phylogeny including 107 high-quality genomes of 107 birds (right). Species names are provided for genomes with a gene loss. High-quality genomes were chosen based on the abundance of inactivation mutations. Genomes with inactivating mutations in >1,900 genes were excluded from the analysis.

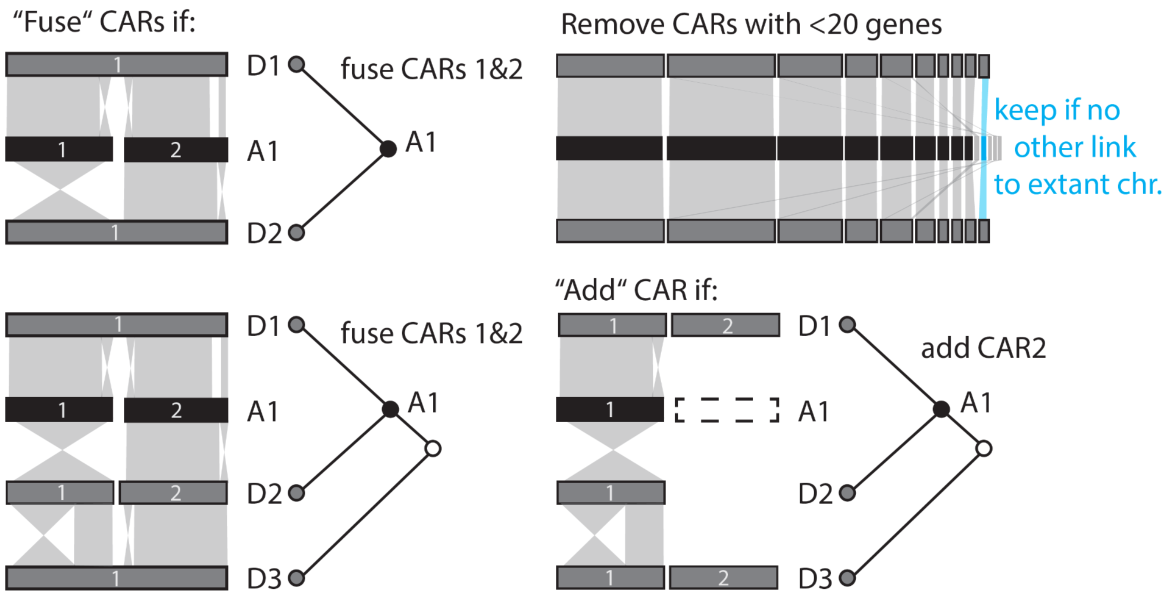


**Figure S13. Manual curation of ancestral chromosome evolution.** For each type of manual curation, illustrations show changes in genome architecture across reconstructed ancestors (black, A1) and descendents (grey, D1-3) (left) together with their phylogenetic relationships (right). We corrected over splitting of ancestral chromosomes by fusing CARs if they were adjacent in both descendent lineages, we re-named them and treated them as a single ancestral chromosome in subsequent analyses (CAR_X, CAR_Y → CAR_XA CAR_XB). Similarly, CARs were fused, if they were adjacent in one descending lineage and in the most closely related outgroup. To focus on chromosome-size CARs, we removed CARS containing less than 20 genes, unless they were the only representative of a chromosome in a descending lineage. Finally, to account for potentially missing chromosome size representation of chromosomes in assemblies of extant species, we assumed a CAR was falsely missing in an ancestor, if it was present in one descending lineage and the most closely related outgroup. We started the curation at genome assemblies of extant species followed by increasingly older ancestral nodes so that already curated genomes would serve as references for further curation of their ancestors.

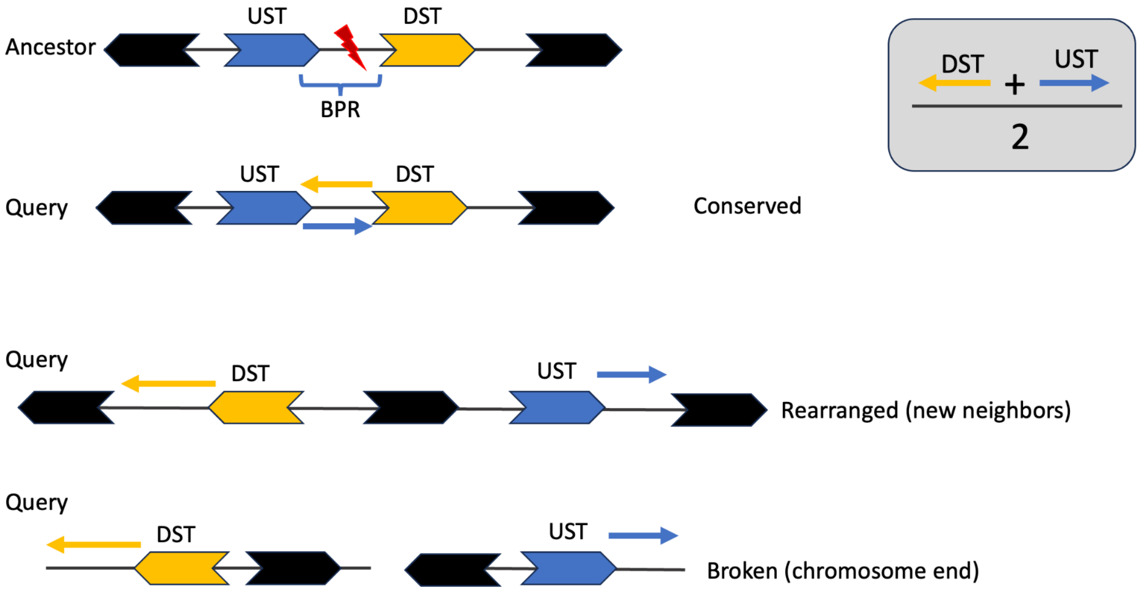


**Figure S14. Approach to extract breakpoint region features in extant query genomes.** From top to bottom, this figure illustrates the breakpoint (red lightning) region in the last ancestor in which the region was not fissioned (top), a rearranged query, and a fissioned query. To extract breakpoint window features in potentially rearranged genomes, we filtered genome annotations of extant species to include only genes (thick) present in the most recent reconstructed ancestor before the fission occurred. Using these filtered annotations, we extracted breakpoint region features for subsequent analyses in extant query genomes by i) selecting the region between the upstream breakpoint gene (UST) and its next downstream neighboring gene (DST), ii) selecting the region between the downstream breakpoint gene and its next upstream neighboring gene and iii) calculating the average repeat content of i) and ii) (grey box). If the upstream/downstream breakpoint gene was the last/first on a chromosome, we considered the region between that gene and the end/start of the scaffold.
